# CD55–CD319–CX3CR1 flow cytometry gating strategy recapitulates scRNA-seq–defined memory CD8 T cell subpopulations

**DOI:** 10.64898/2026.09.15.751807

**Authors:** Pavla Bohacova, Marina Terekhova, Oleg Shpynov, Thomas Francis, Kamila Husarcikova, Petr Tsurinov, Jan Kossl, Maksim Kleverov, Molly Keppel, Stephen D.R. Harridge, Nathan Singh, Maxim N. Artyomov

**Affiliations:** Department of Pathology and Immunology, Washington University School of Medicine, Saint Louis, MO, USA; JetBrains Research, Munich, Germany; Centre for Human and Applied Physiological Sciences, School of Basic and Medical Biosciences, Faculty of Life Sciences & Medicine, King’s College London, London, SE1 1UL, UK; JetBrains Research, Paphos, Cyprus; Division of Oncology, Department of Medicine, Washington University School of Medicine, St. Louis, MO, USA; Center for Gene and Cellular Immunotherapy; Division of Oncology, Section of Cellular Therapies, Washington University School of Medicine

**Keywords:** human CD8 T cells, immunophenotyping, flow cytometry, scRNA-seq, gating strategy, T-cell heterogeneity

## Abstract

Human CD8 T cells have traditionally been classified into naive, central memory, effector memory, and terminal effector subsets using CCR7 and CD45RA expression, a framework that has guided immunological research and clinical immune monitoring for nearly three decades. However, recent single-cell studies have revealed transcriptionally distinct CD8 T cell populations, including GZMK-, GZMB-, and central memory-like states, raising important questions regarding their relationship to canonical flow cytometric subsets. Here, we systematically integrated transcriptomic, epigenetic, and phenotypic analyses to evaluate the correspondence between these classification schemes. We demonstrate that conventional CCR7–CD45RA gating generates heterogeneous populations containing extensive mixtures of transcriptionally and epigenetically distinct CD8 T cell states, resulting in poor resolution of biologically meaningful subsets. To address this limitation, we developed a surface-marker framework based on CD55, CD319, and CX3CR1 that accurately identifies transcriptionally defined human CD8 T cell populations using standard flow cytometry. This strategy enables direct isolation of viable cells, including GZMK-expressing cells increasingly implicated in aging, chronic inflammation, autoimmunity, and cancer, which previously could only be identified using intracellular staining or single-cell sequencing. Functional characterization of purified subsets revealed marked differences in proliferative capacity, cytokine production, and cytotoxic activity, demonstrating that transcriptionally defined states possess distinct immune functions. Together, these findings establish a biologically grounded framework for CD8 T cell classification and provide a practical platform for mechanistic studies, biomarker discovery, and cellular immunotherapy applications.

## INTRODUCTION

Historically, human CD8 T cells were divided into four major subsets based on the expression of CCR7 and CD45RA, a framework introduced in the late 1990s that became widely adopted across immunology and clinical research^1^. In this canonical model, naive, central memory, effector memory, and terminal effector populations are identified using quadrant gating of CCR7 and CD45RA expression. This classification was originally designed to approximate differences in T cell homing behavior and differentiation state using surface markers associated with lymphoid trafficking and antigen experience^2–7^. Additional CD8 T cell gating strategies, such as CD27–CD45RA and CD62L–CD45RA, have also been used in the field, and they generate subsets broadly analogous to the naive and effector memory subsets defined by canonical CCR7–CD45RA gating^8–12^. Together, these approaches have become a cornerstone of human CD8 T cell classification for nearly three decades, significantly shaping immune monitoring and biomarker discovery in human studies, as well as the translation of T cell biology into clinical practice.

In parallel, wide adoption of the single-cell RNA-seq approaches have dissected blood CD8 T cells compartment using the unbiased clustering approaches. Presently, single-cell studies demonstrated that human CD8 T cells robustly separate into transcriptionally defined subpopulations, including naive cells and memory subpopulations characterized by high expression of GzmK, GzmB, or relatively low granzyme expression corresponding most closely to central memory cells^13–17^. Importantly, a growing body of literature underscores clinical relevance of such subsets. For instance, GZMK-expressing CD8 T cells are enriched across a wide range conditions, including autoimmune diseases, chronic inflammation, and cancer^18–21^, highlighting the need to better understand their biology and function. However, identification of these cells currently relies on either single-cell transcriptomics or intracellular GZMK staining, approaches that do not permit recovery of viable cells. Consequently, the field lacks a practical strategy for isolation of GZMK-expressing CD8 T cells and their direct functional and mechanistic characterization.

Given two parallel and widespread approaches to characterize circulating CD8 T cells, important question arises: how do transcriptionally defined CD8 T cell subpopulations relate to the canonical flow cytometry-defined subsets^22^? Here, we demonstrate that canonical CCR7–CD45RA gating yields low-purity populations, resulting in extensive mixing of transcriptionally and epigenetically distinct cell populations within canonical subsets. This disconnect has major implications for both research and clinical applications, as assays performed on heterogeneous mixtures of cellular states may fail to accurately link molecular identity with biological function^22^. Reliance on canonical gating strategies may therefore obscure biologically important differences between CD8 T cell populations and limit the ability to accurately identify, isolate, and optimize cell states with desirable functional properties for adoptive cellular therapies.

To overcome these limitations, we developed an alternative flow cytometry gating strategy based on combination of CD55–CD319–CX3CR1 surface markers that accurately recapitulates transcriptionally and epigenetically defined human CD8 T cell subpopulations. Unlike transcriptomics-based classifications, this approach enables rapid identification, viable cell sorting, and downstream functional assays. Using this framework, we demonstrate that isolated CD8 T cell populations exhibit profoundly divergent proliferative, cytokine-producing, and cytotoxic properties. Together, these findings establish the CD55–CD319–CX3CR1 framework as a biologically accurate strategy for characterization and isolation of primary human CD8 T cell populations in both research and clinical settings.

## RESULTS

### Canonical flow cytometry gating subsets do not match scRNA-seq–defined CD8 T cell subpopulations

To evaluate how canonical flow cytometry-defined CD8 T cell subsets correspond to transcriptionally defined cell subpopulations (**Fig. 1A**), we performed scRNA-sequencing on individually sorted canonical CD8 T cell subsets defined by conventional CCR7–CD45RA gating. Fluorescence minus one (FMO) controls were used to ensure accurate gate placement (**Fig. S1A, B**). Pooled samples were integrated by donor-level and by the guidance of unsupervised clustering we identified four major transcriptionally distinct clusters (**Fig. 1B, S1C, Table1**). Consistent with previous reports^13,15,23,24^ (**Fig. S2**), these comprised naive cells and three major memory populations: central memory cells (Tcm; granzyme-low subset), GZMK-expressing memory cells (TemK), and GZMB-expressing memory cells (TemB) (**Fig. 1B, C, Table1**). A small KLRC2^+^ Tmem population was also detected; however, owing to its very low frequency (∼0.5% of CD8 T cells in adults^13^), it was not investigated further in this study. Notably, canonical CCR7–CD45RA-defined memory subsets were broadly dispersed across the transcriptional UMAP and exhibited extensive overlap with other sorted subsets (**Fig. 1B**), suggesting that conventional gating does not faithfully reflect the transcriptional subsets underlying human CD8 T cell heterogeneity.

**Figure 1.**
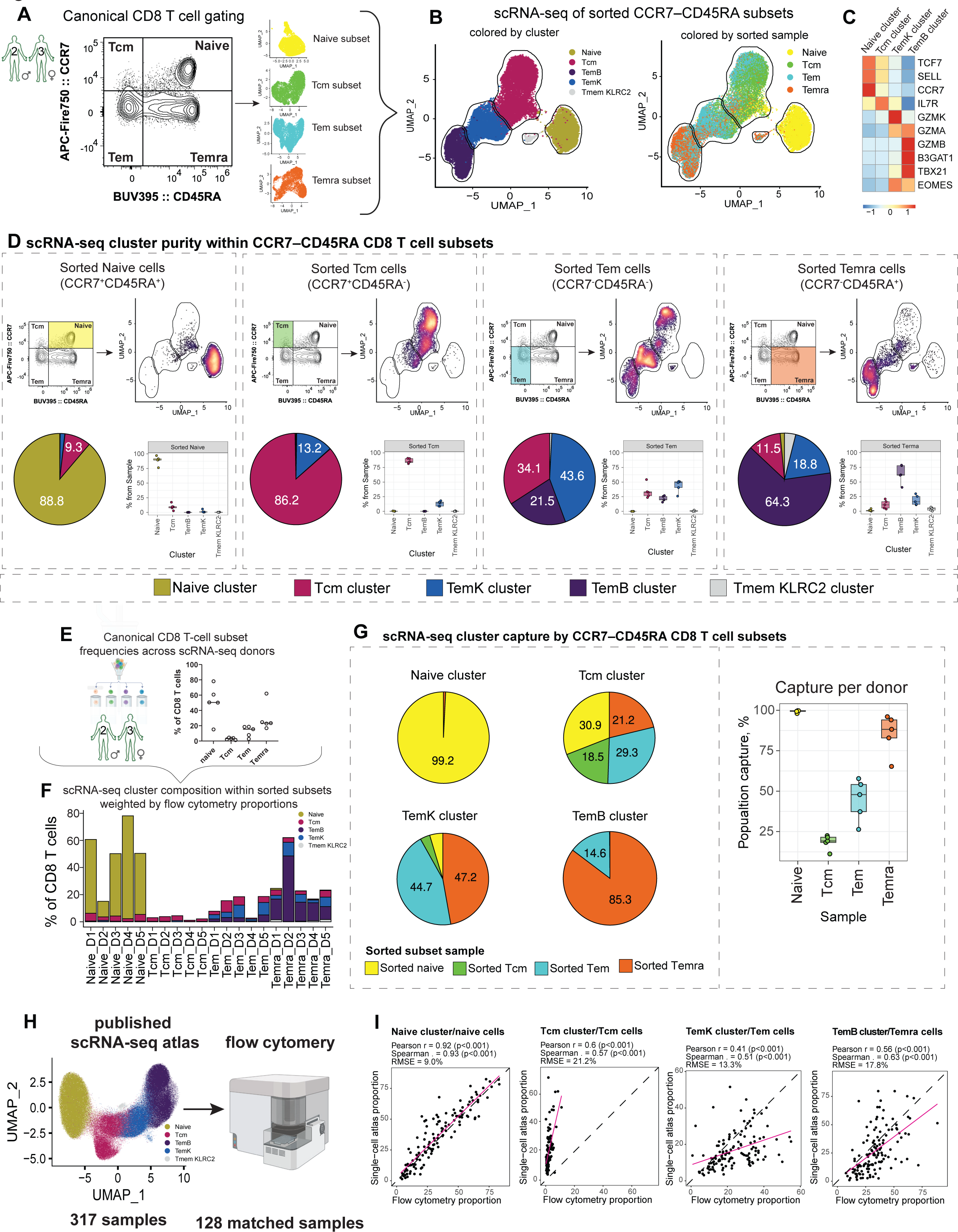
Transcriptional heterogeneity within canonical CD8 T cell subsets. (A) Representative flow cytometry plot of canonical CCR7–CD45RA gating. (B) Uniform manifold approximation and projection (UMAP) plot of CCR7–CD45RA subsets, colored by cluster (left), colored by sorted subsets (right). Borders correspond to transcriptional clusters (C) Heatmap showing normalized and scaled expression of selected genes across clusters. (D) Analysis of transcriptional cluster purity within sorted CCR7–CD45RA subsets. Representative gating strategy (top left), UMAP density plots with borders correspond to transcriptional clusters (top right), cluster composition pie charts (bottom left), and box plots showing the percentage of cells of each CD8 T cell cluster across donors (bottom right). (E) Scatter plot showing proportions of canonical CD8 T cell subsets across donors used for scRNA-seq analysis of CCR7–CD45RA subsets. (F) Bar plot summarizing transcriptional composition of canonical CD8 T cell subsets in relation to flow cytometry subset proportions for donors used for scRNA-seq analysis of CCR7–CD45RA subsets. (G) Analysis of scRNA-seq cluster capture by CCR7–CD45RA subsets. Pie charts showing distribution of canonical gates capturing each transcriptionally defined population (left), box plots showing the percentage of cells from each scRNA-seq cluster captured by individual CCR7– CD45RA gates across donors (right). (H) Schematic overview of direct comparison of matched samples from the ABF300 cohort. (I) Correlation of canonical flow cytometry subset frequencies with matched scRNA-seq-defined populations across donors. Correlation coefficients and RMSE values are indicated. Dashed line indicates the line of identity (y = x); pink line indicates the linear regression fit. In (D) and (G), box plot hinges indicate 25th and 75th percentiles; whiskers extend to 1.5×IQR; horizontal bars denote medians.

Therefore, we proceeded to quantify the degree of this inconsistency. While sorted CCR7^+^CD45RA^+^ naive and CCR7^+^CD45RA^−^ Tcm populations consisted of predominantly their scRNA-seq-defined counterparts (∼88% Naive and ∼86% Tcm cluster cells, respectively) (**Fig. 1D**), CCR7^−^CD45RA^−^ Tem and CCR7^−^CD45RA^+^ Temra populations consisted of mixtures of transcriptional heterogeneous subpopulations (**Fig. 1D**). Notably, the canonical Tem gate lacked a dominant scRNA-seq-defined counterpart, with cells distributed across Tcm (∼21%), TemK (∼43%), and TemB (∼34%) clusters, whereas the Temra gate remained substantially contaminated by non-TemB populations, including TemK (∼18%) and Tcm (∼11%) cluster cells (**Fig. 1D**). These results demonstrate that canonical gating yields highly mixed, low-purity effector memory CD8 T-cell populations when evaluated against scRNA-seq-defined cell subpopulations. We next evaluated how efficiently canonical CCR7–CD45RA gating captured the corresponding scRNA-seq-defined transcriptional populations (**Fig. 1E–G**). Specifically, we asked what fraction of cells belonging to a given transcriptional cluster, such as the central memory cluster, would be identified within the corresponding canonical CCR7⁺CD45RA⁻ gate. To quantify this, we considered the frequency of each canonical flow cytometry gate (**Fig. 1E**) and its transcriptional composition (**Fig. 1F**), allowing us to estimate the fraction of each scRNA-seq-defined population captured by canonical gating. Using this approach, we found that canonical gating strategy failed to capture the majority of scRNA-seq-defined central memory CD8 T cells. On average, less than 20% of all Tcm cluster cells were contained within the canonical CCR7^+^CD45RA^−^ Tcm gate, whereas the remaining cells were distributed across other subsets (**Fig. 1G**). Similarly, less than 45% of TemK cluster cells were located within the corresponding canonical Tem gate. These results indicate that canonical CCR7–CD45RA gating also fails to efficiently capture transcriptionally defined CD8 T cell populations within the corresponding canonical subsets.

Lastly, we directly compared the frequencies of CD8 T cell populations defined by canonical CCR7–CD45RA gating and scRNA-seq on the large cohort level. To this end, we immunophenotyped 128 samples matched to the samples used in ABF300 scRNA-seq dataset^13^ (**Fig. 1H**). Correlation analysis demonstrated that naive cell frequencies showed strong agreement between canonical gating and scRNA-seq-defined populations, whereas memory subsets displayed substantially weaker concordance (**Fig. 1I, Table1**), especially in the context of central memory cells. Together, these findings demonstrate that canonical CCR7–CD45RA gating poorly reflects scRNA-seq-defined memory CD8 T cell subpopulations.

### CD55–CD319–CX3CR1 flow cytometry gating strategy achieves high transcriptional purity and capture of scRNA-seq-defined CD8 T cell populations

Based on these observations, we sought to develop a novel surface marker-based gating strategy for conventional CD8 T cells that would align with scRNA-seq-defined memory cell subpopulations while remaining compatible with viable cell sorting and downstream functional assays. Naive cells were identified using FAS–CCR7 gating, as described before^14^. To identify candidate markers for memory populations, we generated pseudobulk expression profiles from the scRNA-seq data shown in Fig. 1B (**Fig. 2A**) and first noted that key transcriptional populations can be distinguished by levels of granzymes expression. The cluster of central memory cells (Tcm) was characterized by low level of granzymes expressions. Effector memory cells highly expressed GzmA and GzmM transcripts and could be distinguished between each other by high levels of GzmK or GzmB expression correspondingly (TemK and TemB). To evaluate candidate gating strategies, we first assessed granzyme expression levels within the corresponding gates, then evaluated transcriptional enrichment using bulk RNA-seq of sorted populations, and finally performed single-cell RNA-seq of sorted subsets as the gold-standard validation (**Fig. 2A**).

**Figure 2.**
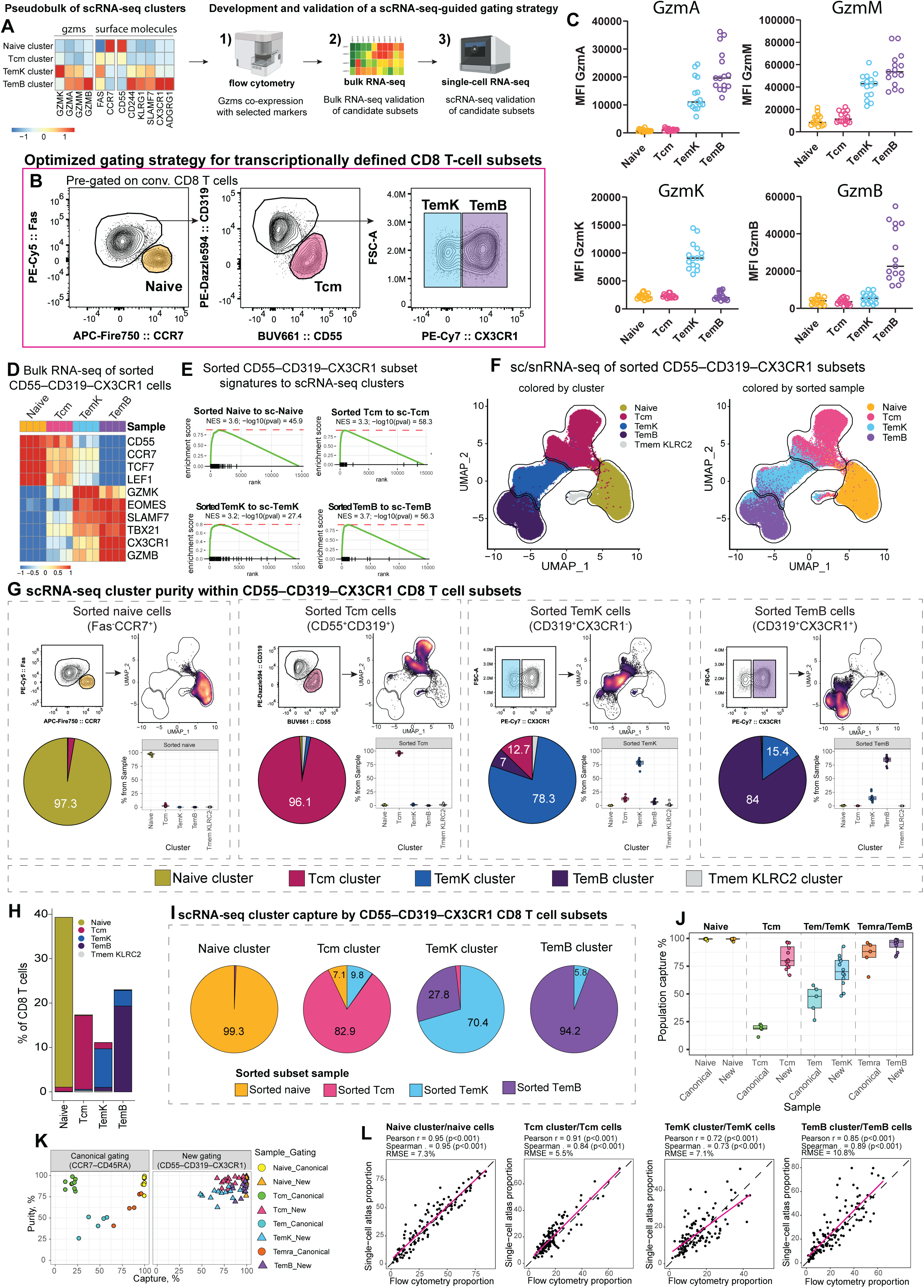
CD55–CD319–CX3CR1 gating resolves transcriptional CD8 T cell clusters. (A) Schematic overview of development of the CD55–CD319–CX3CR1 gating strategy. On left, heatmap showing normalized and scaled expression of selected genes across clusters For data shown on Fig 1B. (B) Representative flow cytometry plots showing gating strategy defining Naive, Tcm, TemK, and TemB CD8 T cell subsets. (C) Scatter plots showing median fluorescence intensity (MFI) for protein expression of granzymes in CD55–CD319–CX3CR1 subsets. (D) Heatmap showing normalized and scaled expression of selected genes of sorted CD55– CD319–CX3CR1 subsets. (E) Gene set enrichment analysis (GSEA) plot of sorted CD55–CD319–CX3CR1 subsets enriched to scRNA-seq of CCR7–CD45RA subsets from Fig.1B. (F) UMAP plot of sorted CD55–CD319–CX3CR1 subsets, colored by cluster (left), colored by sorted subsets (right). Borders correspond to transcriptional clusters. (G) Analysis of transcriptional cluster purity within sorted CD55–CD319–CX3CR1 subsets from sc/snRNA-seq. Representative gating strategy (top left), UMAP density plots with borders correspond to transcriptional clusters (top right), cluster composition pie charts (bottom left), and box plots showing the percentage of cells of each CD8 T cell cluster across donors (bottom right). (H) Bar plot summarizing transcriptional composition CD55–CD319–CX3CR1 subsets in relation to flow cytometry subset proportions used for scRNA-seq analysis of CD55–CD319–CX3CR1 subsets (mean of all donors). (I) Analysis of sc/snRNA-seq cluster capture by CD55–CD319–CX3CR1 subsets. Pie charts showing distribution of canonical gates capturing each transcriptionally defined population. (J) Box plots showing the percentage of cells from each sc/snRNA-seq cluster captured by individual CCR7–CD45RA or CD55–CD319–CX3CR1 gate across donors. (K) Scatter plots showing the percentage of cells from each sc/snRNA-seq cluster captured by individual CCR7–CD45RA or CD55–CD319–CX3CR1 gate across donors. (L) Correlation of CD55–CD319–CX3CR1 flow cytometry subset frequencies with matched scRNA-seq-defined populations across donors. Correlation coefficients and RMSE values are indicated. Dashed line indicates the line of identity (y = x); pink line indicates the linear regression fit. In (G) and (J), box plot hinges indicate 25th and 75th percentiles; whiskers extend to 1.5×IQR; horizontal bars denote medians.

We closely evaluated CD55, KLRG1, CD244, CD319, CX3CR1 and gpr56 (**Fig S3, Table1**). The combination of CD55 and CD319 provided the clearest separation and captured a larger fraction of GzmA- and GzmM-expressing cells (**Fig. S3A-D**), identifying these markers as promising discriminators between Tcm and Tem populations. For discrimination of TemK and TemB populations, we closely evaluated both CX3CR1 and gpr6 based on their published association with GZMB expression^25–28^. As both markers showed comparable performance and highly overlapping expression (**Fig. S3F-G**), CX3CR1 was selected because of the broader availability of conjugated antibodies. Collectively, these analyses suggested that CD55–CD319–CX3CR1 gating strategy could isolate CD8 T cell subsets corresponding to scRNA-seq-defined populations (**Fig 2B, C, S4A, B**), in contrast to conventional gating strategies (**Fig. S4C, Table1**).

To further benchmark the CD55–CD319–CX3CR1 gating strategy, we sorted CD55–CD319– CX3CR1-defined CD8 T cell populations (FMO controls guided gate placement) (**Fig. S5A, B**) and performed bulk RNA-seq to determine whether these subsets are transcriptionally distinct (**Fig. 2D, S5C, Table1**). Gene set enrichment analysis confirmed that the sorted subsets corresponded to their expected scRNA-seq-defined cluster identities (**Fig. 2E**). As ultimate proof, we analyzed gene expression data from scRNA-seq and multiome sequencing of sorted CD55– CD319–CX3CR1-gated CD8 T cells (**Fig. 2F, S5D, Table1**). Combined and integrated data were clustered, yielding Naive, Tcm, TemK, and TemB clusters consistent with clusters from **Fig.1** described above (**Fig. S5E**). Coloring the UMAP by individual sorted populations revealed clear segregation in accordance with the transcriptional clusters (**Fig. 2F**). Sorted CCR7^+^Fas^−^ CD8 T cells (Naïve) and CD55^+^CD319^−^ CD8 T cells (Tcm) populations showed remarkably high concordance with their scRNA-seq-defined counterparts, consisting of ∼97% naïve and ∼96% Tcm cluster cells (**Fig. 2G**). Likewise, CD319^+^CX3CR1^+^ CD8 T cells (TemB) and CD319^+^CX3CR1^−^CD8 T cells (TemK) populations also showed strong enrichment for their corresponding transcriptional states, containing ∼84% and ∼78% of TemB and TemK cluster cells, respectively. Together, these results demonstrate that the CD55–CD319–CX3CR1 gating strategy yield highly pure cell subpopulation in terms of the transcriptional cell subsets.

Next, we evaluated the ability of the CD55–CD319–CX3CR1 gating to efficiently capture scRNA-seq-defined CD8 T cell subsets within their corresponding flow cytometry gates (**Fig. 2H-K**). In striking contrast to canonical CCR7–CD45RA gating, the CD55–CD319–CX3CR1 gating strategy demonstrated high assignment accuracy across all major CD8 T cell populations. Specifically, 99% of Naïve, 82% of Tcm, 70% of TemK, and 94% of TemB cluster cells localized within their corresponding gated subsets (**Fig. 2H, I**), which represents a substantial improvement over canonical CCR7–CD45RA gating (**Fig. 2J, K**). Importantly, when different classical gating strategies (CCR7–CD45RA, CD62L–CD45RA, and CD27–CD45RA) were compared against the CD55–CD319–CX3CR1 framework, all classical approaches showed substantial mixing of transcriptionally distinct CD8 T cell populations defined by CD55–CD319–CX3CR1 gating (**Fig. S6**). Lastly, in large cohort measurements, the CD55–CD319–CX3CR1 gating strategy demonstrated a high degree of concordance with scRNA-seq-defined CD8 T cell populations, substantially improving upon canonical gating (**Fig. 2L** compared to **Fig. 1I**). Together, these findings demonstrate that the CD55–CD319–CX3CR1 gating strategy faithfully recapitulates transcriptionally defined CD8 T cell states across individuals and enables accurate flow cytometry-based quantification of scRNA-seq-defined populations.

### CD55–CD319–CX3CR1-defined CD8 T cell subpopulations are epigenetically distinct

We next asked whether CD55–CD319–CX3CR1-defined CD8 T cell subsets also represent epigenetically distinct populations. As transcriptional profiling data in **Fig.2F** included both single-cell RNA-seq and multiome data, we next analyzed the corresponding chromatin accessibility profiles from the same sorted CD8 T cell subpopulations. UMAP visualization based on snATAC-seq peak accessibility revealed clear segregation of the major transcriptionally defined CD8 T cell subsets (**Fig. 3A, Table1**), suggesting substantial underlying epigenetic differences. Importantly, the sorted populations were also well separated in the single-nuclei ATAC UMAP space (**Fig. 3A, B**), confirming that CD55–CD319–CX3CR1-based gating captures biologically distinct epigenetic states, with each subset exhibiting increased chromatin accessibility at its characteristic genes (**Fig. 3C, D**). Moreover, differential accessibility analysis identified many uniquely accessible chromatin regions, further supporting their epigenetic distinctness (**Fig. 3E**). Shared regulatory programs were observed primarily between Naive and Tcm cells, as well as between TemK and TemB cells, which may reflect their relationship along the CD8 T cell differentiation trajectory and shared functional programs.

**Figure 3.**
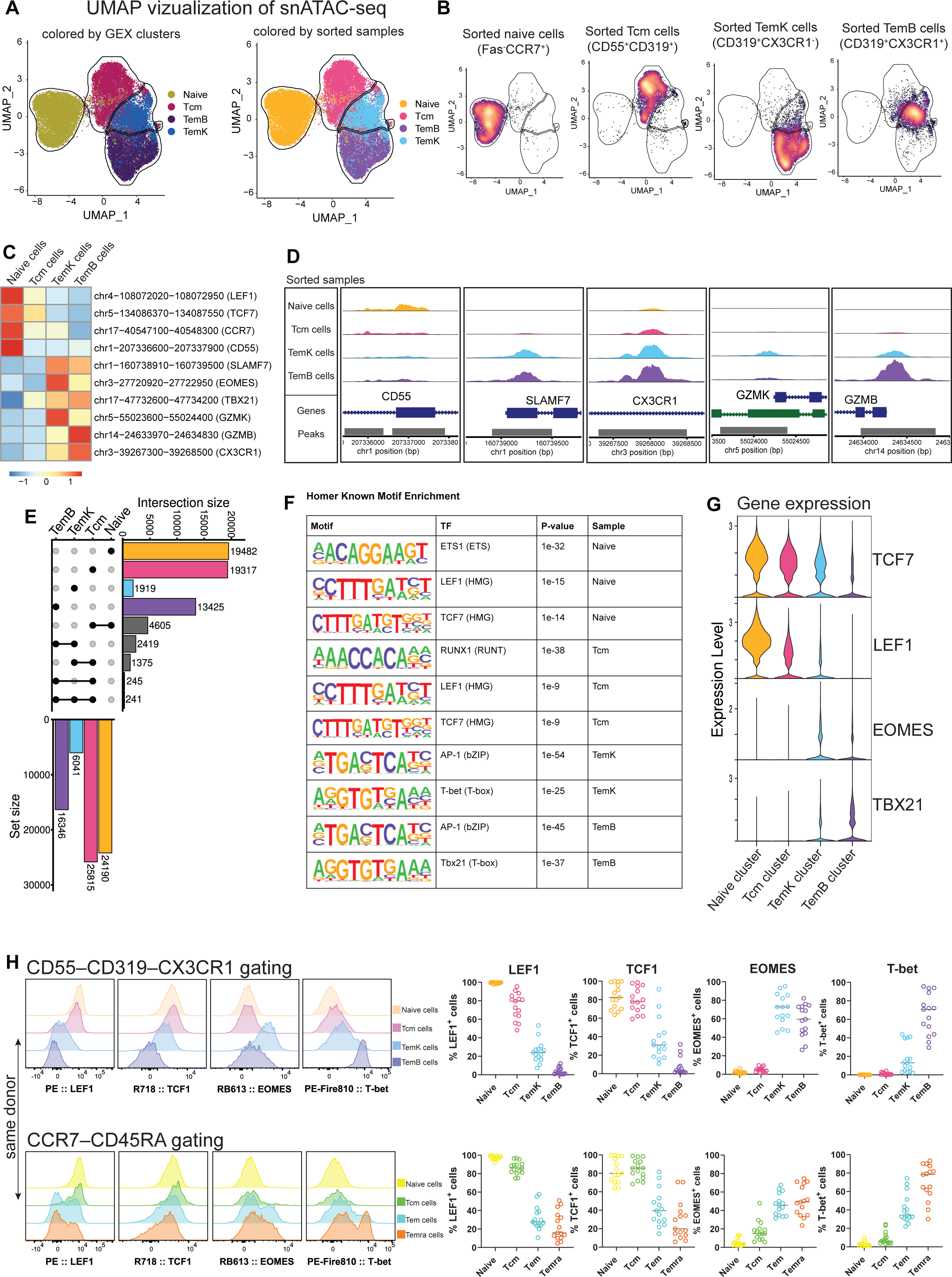
Single-nuclei analysis of chromatin accessibility of sorted CD55–CD319–CX3CR1 subsets. (A) UMAP plot based on snATAC-seq of CD55–CD319–CX3CR1 subsets, colored by transcriptional cluster defined in Fig. 2F (left), colored by sorted subsets (right). Borders from transcriptional clusters from Fig. 2F. (B) snATAC-seq UMAP density plots with borders correspond to transcriptional clusters from Fig. 2F. (C) Heatmap showing TF-IDF-normalized and scaled chromatin accessibility of selected ATAC peaks for CD55–CD319–CX3CR1 subsets. (D) Coverage plots showing peaks of accessible chromatin near selected genes. (E) UpSet plot showing the overlap of differentially accessible regions (DARs) between CD55– CD319–CX3CR1 subsets. Only DARs with adjusted P < 1 × 10⁻⁵ were included. Only intersections containing more than 50 regions are shown. (F) Established transcription factor motif analysis (“Known motifs”) of CD55–CD319–CX3CR1 subset-specific regions. (G) Violin plots showing normalized expression of selected transcription factor genes per sample from Fig.2F. (H) Representative histograms showing protein transcription factor expression by CD55– CD319–CX3CR1 or CCR7–CD45RA subsets (left). Scatter plots showing the percentage of transcription factor positive cells CD55–CD319–CX3CR1 or CCR7–CD45RA subsets (right).

To identify potential regulators of these states, we performed motif enrichment analysis. This revealed strong enrichment of LEF1 and TCF7 motifs in Naive and Tcm cells, whereas AP-1 family and T-box-family motifs were preferentially enriched in TemK and TemB populations (**Fig. 3F**). Notably, these findings complemented both the accessibility of the corresponding transcription factor loci and their expression patterns observed in the scRNA-seq data (**Fig. 3G**). Together, these results indicate that LEF1, TCF7, EOMES, and TBX21 are not only expressed and epigenetically accessible, but are also likely active regulators contributing to the establishment of distinct CD8 T cell states. Moreover, CD55–CD319–CX3CR1-defined subsets were selectively enriched for cells expressing LEF1, TCF1 (encoded by TCF7), EOMES, and T-bet (encoded by TBX21) at the protein level (**Fig. 3H, S7A**), whereas conventional gating strategies showed substantial overlap of cells expressing these transcription factors across subsets (**Fig. 3H, Fig. S7B, C, Table1**). Collectively, these findings demonstrate that CD55–CD319–CX3CR1-based gating identifies epigenetically distinct CD8 T cell subpopulations.

### CD55–CD319–CX3CR1-defined CD8 T cell subpopulations exhibit distinct effector functions

Given the distinct transcriptional and epigenetic identities of CD55–CD319–CX3CR1-defined CD8 T cell subpopulations, we next investigated whether these subpopulations also differ functionally. Sorted subsets were stimulated with αCD3/αCD28 beads and evaluated for activation dynamics and cytokine production (**Fig. 4A**). Following stimulation, all subsets maintained viability throughout the time course (**Fig. S8A, B, Table1**). However, their proliferative responses diverged substantially. Fas^-^CCR7^+^ naive and CD55^+^CD319^-^ Tcm populations underwent robust expansion beginning at day 2, whereas proliferation of CD319^+^CX3CR1^-^ TemK cells was slower and delayed. In contrast, CD319^+^CX3CR1^+^ TemB cells displayed minimal proliferative capacity and the population declined over the course of stimulation (**Fig. 4B, S8C, Table1**). These findings highlight a key limitation of bulk CD8 T cell functional studies and *ex vivo* expansion approaches such as CAR-T manufacturing, as differential subset proliferation can rapidly alter culture composition relative to the original cell subset distribution.

**Figure 4.**
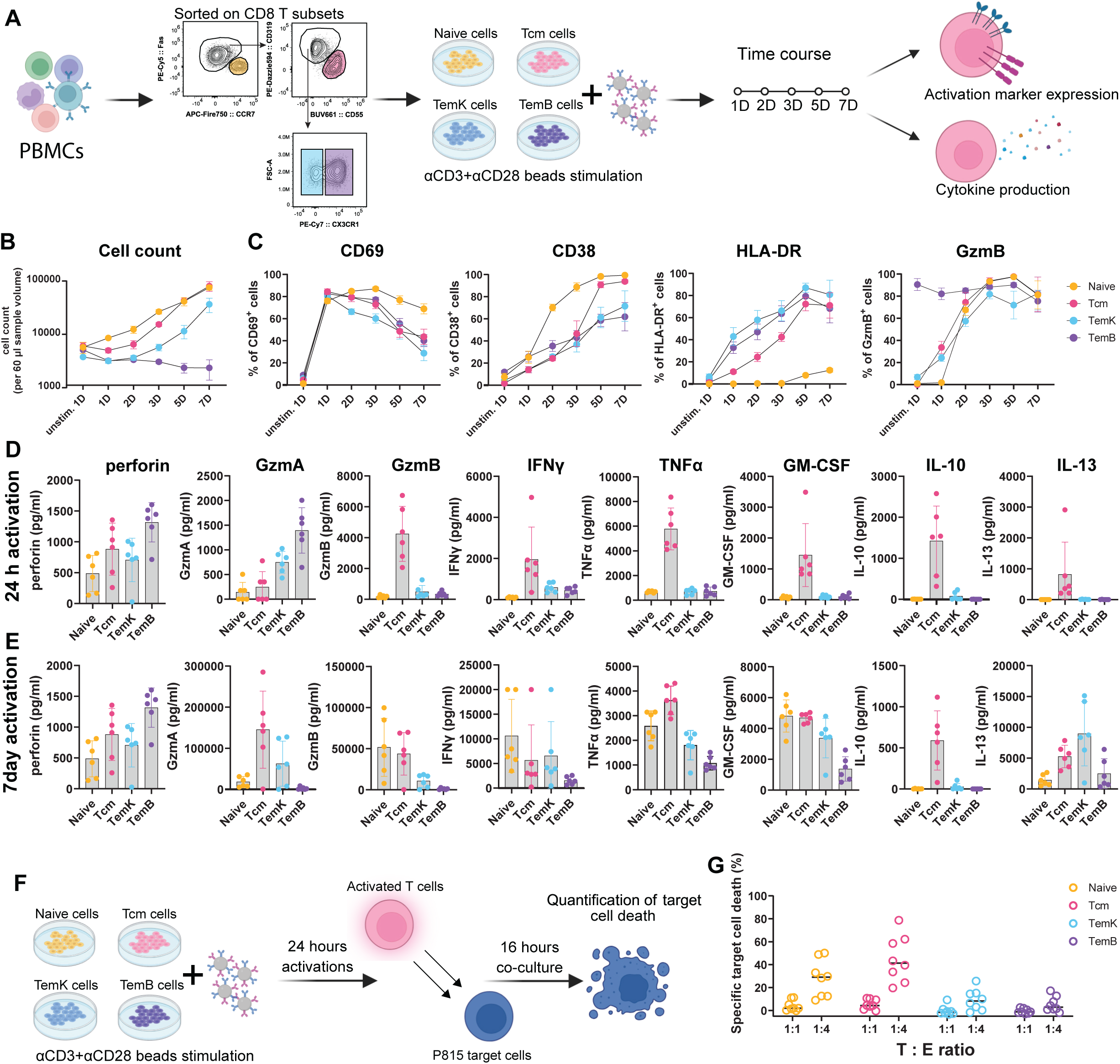
Functional responses of CD55–CD319–CX3CR1-defined subsets. (A) Overview illustrating the functional assays of CD8 T subsets. (B) Line plots showing cell expansion kinetics of sorted CD55–CD319–CX3CR1 subsets, logarithmic scale. (C) Line plots showing expression kinetics of selected protein markers by sorted CD55–CD319– CX3CR1 subsets. (D) Bar plots showing cytokine production upon 24 hours after αCD3/αCD28 bead activation. (E) Bar plots showing cytokine production after 7 days of αCD3/αCD28 bead activation. (F) Schematic overview of redirected cytotoxicity assay. (G) Scatter plot quantifying specific target cell death induced by activated CD55–CD319–CX3CR1 subsets at indicated effector-to-target (E:T) ratios.

To dissect activation-associated functional programs across CD55–CD319–CX3CR1-defined CD8 T cell subsets, we monitored expression of the activation markers CD69, CD38, and HLA-DR and cytotoxic GzmB molecule over the course of stimulation (**Fig. 4C**). As expected, CD69, as an early activation marker, was rapidly induced by all subsets, whereas CD38 and HLA-DR increased progressively throughout the stimulation period and peaked at later stages of activation^29–31^. The most striking difference was the near absence of HLA-DR expression in naive CD8 T cells. In parallel, all subsets rapidly acquired intracellular GzmB expression following activation and became largely GzmB-positive by day 2. TemB cells already exhibited high constitutive GzmB expression prior to stimulation, consistent with their pre-existing cytotoxic phenotype.

We further investigated secretion of effector molecules upon activation of individual CD8 T cell subsets. To this end, we quantified production of a broad panel of cytokines and cytotoxic molecules following short-term (24 h) and prolonged (7 d) stimulation (**Fig. 4D, S8D**). Strikingly, CD55^+^CD319^-^ Tcm cells emerged as the dominant early producers of soluble effector molecules during the first 24 hours of activation, exhibiting broad polyfunctional cytokine secretion of molecules including GZMB, IFNγ, TNFα, GM-CSF, IL-10 and IL-13. Production of GzmA, perforin, and CCL5 was highest by CD319^+^CX3CR1^+^ TemB cells. Prolonged stimulation substantially altered the functional landscape of activated CD8 T cell subsets. After 7 days, naive cells also acquired the capacity to produce multiple cytokines, converging toward the highly polyfunctional phenotype. CD319^+^CX3CR1^-^ TemK likewise gained ability to produce cytokines after long stimulation. Nevertheless, important subset-specific differences persisted. Naive cells remained largely incapable of producing GzmA and Th2-associated cytokines, including IL-4, IL-5, IL-10, and IL-13, whereas TemK cells showed minimal induction of IL-4 and IL-10 despite prolonged stimulation.

Building on these observations, we asked whether CD55–CD319–CX3CR1-defined CD8 T cell subsets also differed in their direct cytotoxic capacity using an *in vitro* redirected killing assay (**Fig. 4F**). Surprisingly, Tcm cells exhibited the strongest cytotoxic activity, reaching the highest levels of specific target cell death (**Fig. 4G, S8E, Table1**). Naive cells also demonstrated clear killing capacity. In contrast, TemK and TemB subsets displayed only limited cytotoxic activity under these assay conditions. Importantly, because CD8 T cell subsets were activated for only 24 hours prior to co-culture with target cells, these differences would not be attributed to the subset-specific proliferative expansion observed at later stages of activation (**Fig. S8F**). Furthermore, the activation protocol relied on αCD3/αCD28 bead stimulation, whereas TemB cells lacked CD28 expression among the analyzed subsets (**Fig. S8G**). This may have limited their responsiveness to stimulation and contributed to their comparatively weak killing capacity in this assay despite high expression of cytotoxic effector molecules. Collectively, these results demonstrate that CD55–CD319–CX3CR1-defined CD8 T cell subsets are not merely molecularly distinct, but represent functionally specialized populations with profoundly different activation, effector, and cytotoxic properties.

## DISSCUSSION

Recent advances in single-cell transcriptomic and epigenomic profiling have substantially expanded our understanding of human CD8 T cell heterogeneity. For example, these approaches have led to identification of GZMK-expressing CD8 T cells as they seem to be biologically and a clinically important population implicated in aging, chronic inflammation, autoimmune diseases, and cancer^13,16,18–21^. Notably, these cells have thus far been identified primarily through transcriptomic approaches. Nevertheless, conventional CCR7–CD45RA or related gating strategies remain overwhelmingly used across both experimental and clinical immunology, including immune monitoring, functional studies, and adoptive cell therapy applications^9,11,32–35^.

Recent efforts within the field have emphasized the limitations of the current T cell framework^22,31^. Our findings experimentally validate these conceptual concerns by demonstrating that canonical CCR7–CD45RA gating incompletely resolves biologically distinct CD8 T cell populations and results in substantial mixing of transcriptionally and epigenetically divergent cell states.

Here, we establish the CD55–CD319–CX3CR1 framework as a practical gating strategy for prospective isolation of transcriptionally and epigenetically defined human CD8 T cell subpopulations in healthy individuals. While the association between CX3CR1 expression and GZMB-expressing CD8 T cells has been described^28,36,37^, however, CD319 expression has only recently been associated with effector-memory CD8 T cells and cytotoxic effector differentiation^38^, and the association of CD55 with resting human central memory CD8 T cells has not, to our knowledge, been previously recognized. Most importantly, these molecules together form a combinatorial framework that enables accurate resolution of transcriptionally defined CD8 T cell subpopulations compared to canonical gating, as demonstrated by the increased transcriptional and epigenetic separation of the resulting subpopulations and their consistent expression of key differentiation-associated transcription factors, including TCF1, LEF1, EOMES, and T-bet. In addition, CD244, KLRG1, and gpr56 displayed expression patterns broadly analogous to those of CD319 and CX3CR1 and may therefore represent practical alternatives for retrospective interrogation of existing datasets or in situations where panel design constraints preclude inclusion of the optimal marker combination. Nevertheless, the CD55–CD319–CX3CR1 marker combination provided a robust and readily implementable framework for prospective isolation of scRNA-seq-defined CD8 T cell subpopulations. Notably, the CD55–CD319–CX3CR1 strategy achieved greater than 96% transcriptional purity while capturing more than 82% of all transcriptionally defined Tcm cells, compared to less than 20% captured by canonical CCR7– CD45RA gating. This distinction is biologically important because, as demonstrated in our functional assays, Tcm cells represented one of the most functionally active populations among circulating human CD8 T cells. These cells exhibited strong cytokine-producing capacity upon activation while simultaneously retaining direct cytotoxic activity against target cells. These findings are consistent with emerging evidence that central memory-like CD8 T cell populations play critical roles in sustaining durable antitumor immune responses and long-term persistence following adoptive cell therapies^39,40^. Our results extend these observations by providing a practical framework for prospective isolation of biologically distinct human CD8 T cell populations using flow cytometry, with important implications for both mechanistic immunology and future cellular immunotherapy strategies. Together, these findings support a transition from historically defined surrogate classifications toward biologically grounded frameworks informed by single-cell multi-omic profiling.

### Study limitations

A limitation of this study is that we did not further investigate gating of the small KLRC2^+^ Tem population because it represented only a minor fraction of conventional CD8 T cells. In addition, our analysis focused on broad transcriptionally defined populations to capture major patterns in CD8 T cells, although each population could be further resolved into subclusters. Further work will be required to resolve finer transcriptional structure of CD8 T cell compartment.

## Supporting information

Donors

Hashtag_antibodies

Flow_cytometry_antibodies

## AUTHOR CONTRIBUTION

These authors contributed equally: Pavla Bohacova and Marina Terekhova.

P.B. and M.N.A. conceived and designed the study. P.B. performed and analyzed experiments.

M.T. performed computational analysis of multimodal data. O.S. and M.Kl. supported computational data processing. T.F. prepared multiome libraries. P.T. performed clustering analysis of flow cytometry data. K.H. and J.K. helped with experimental processing. M.Ke. performed cell sorting. N.S. and S.D.R.H. contributed to data interpretation. P.B. and M.N.A. wrote the manuscript.

## ACKNOWLEDGEMENTS

The work was supported by the Aging Biology Foundation US grant to M.N.A and the Aging Biology Foundation EU grant to S.D.R.H. We thank the Siteman Flow Cytometry Core at Washington University for cell sorting support. We thank the Genome Technology Access Center at the McDonnell Genome Institute at Washington University School of Medicine for assistance with genomic processing. Schematic illustrations were created in <u>BioRender.com</u>.

## SUPPLEMENTAL INFORMATION

**Supplementary Figure 1:**
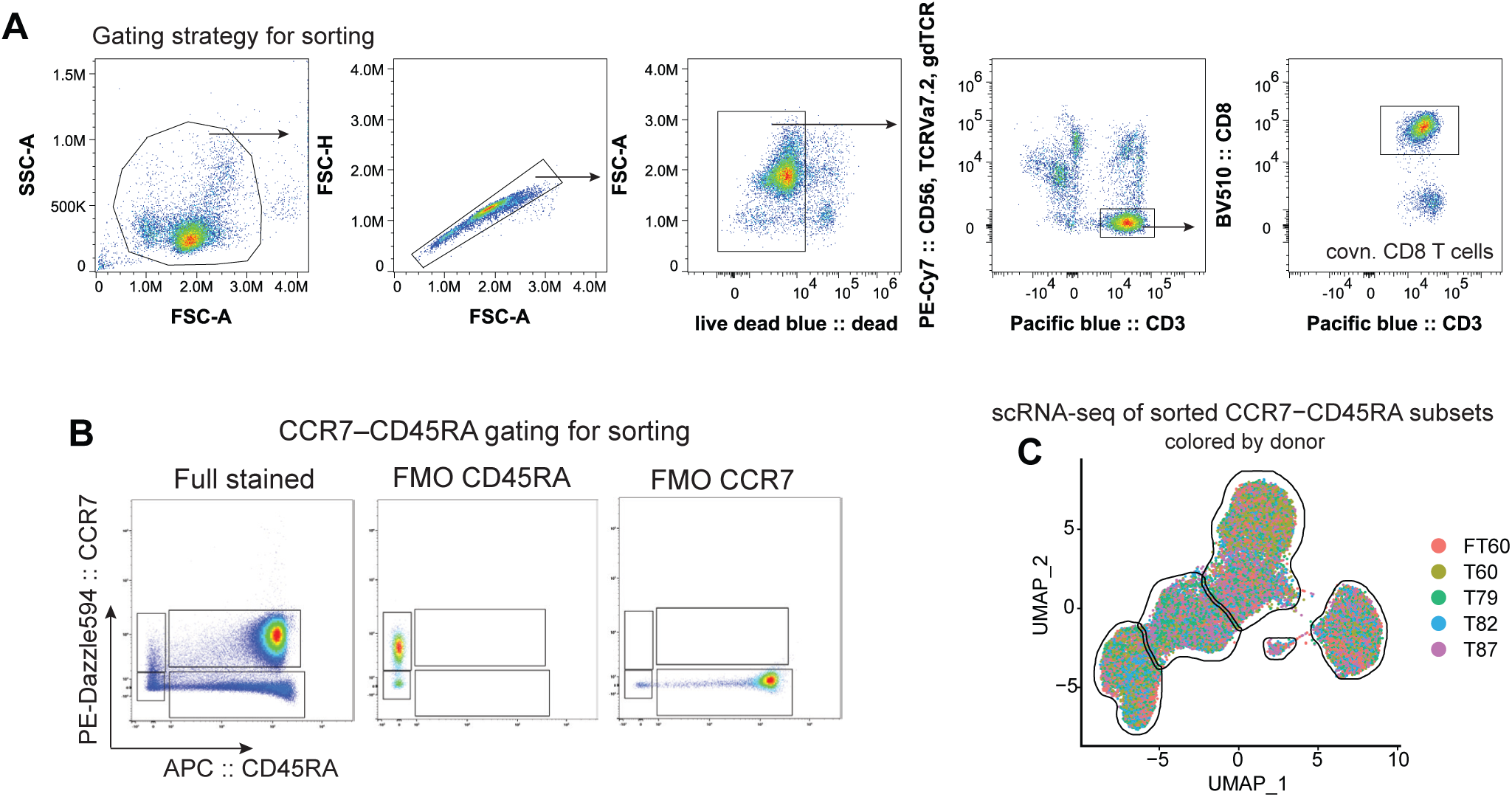
Sorting strategy and scRNA-seq characterization of CCR7– CD45RA subsets related to Figure 1. (A) Representative gating strategy identifying conventional CD8 T cells used for cell sorting. (B) Representative dot plots showing the sorting strategy for CCR7–CD45RA subsets and corresponding FMO controls. (C) UMAP plot of CCR7–CD45RA subsets, colored by donor.

**Supplementary Figure 2:**
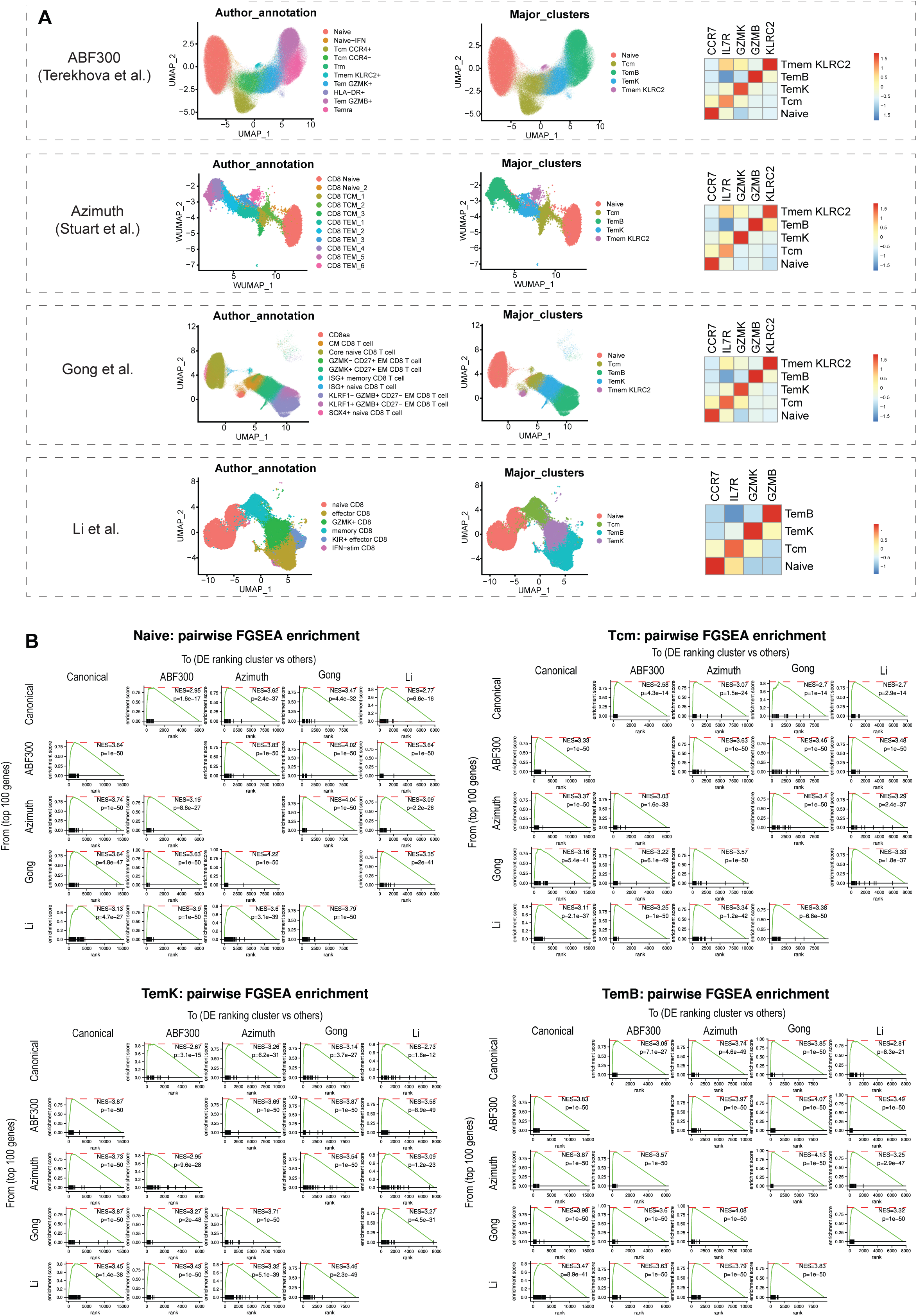
Cross-dataset validation of major human CD8 T cell clusters related to Figure 1. (A) UMAP projections of published human PBMC scRNA-seq datasets showing CD8 T cells, colored by original author annotations (left) or by the four major CD8 T cell clusters (Naive, Tcm, TemK, and TemB) generated by cluster aggregation based on expression of major defining genes. (middle). Heatmaps (right) show normalized and scaled expression of representative marker genes characteristic of the major CD8 T cell clusters. (B) GSEA normalized enrichment score (NES) and p-value for pairwise signature enrichment from selected datasets to corresponding major CD8 T cell clusters.

**Supplementary Figure 3:**
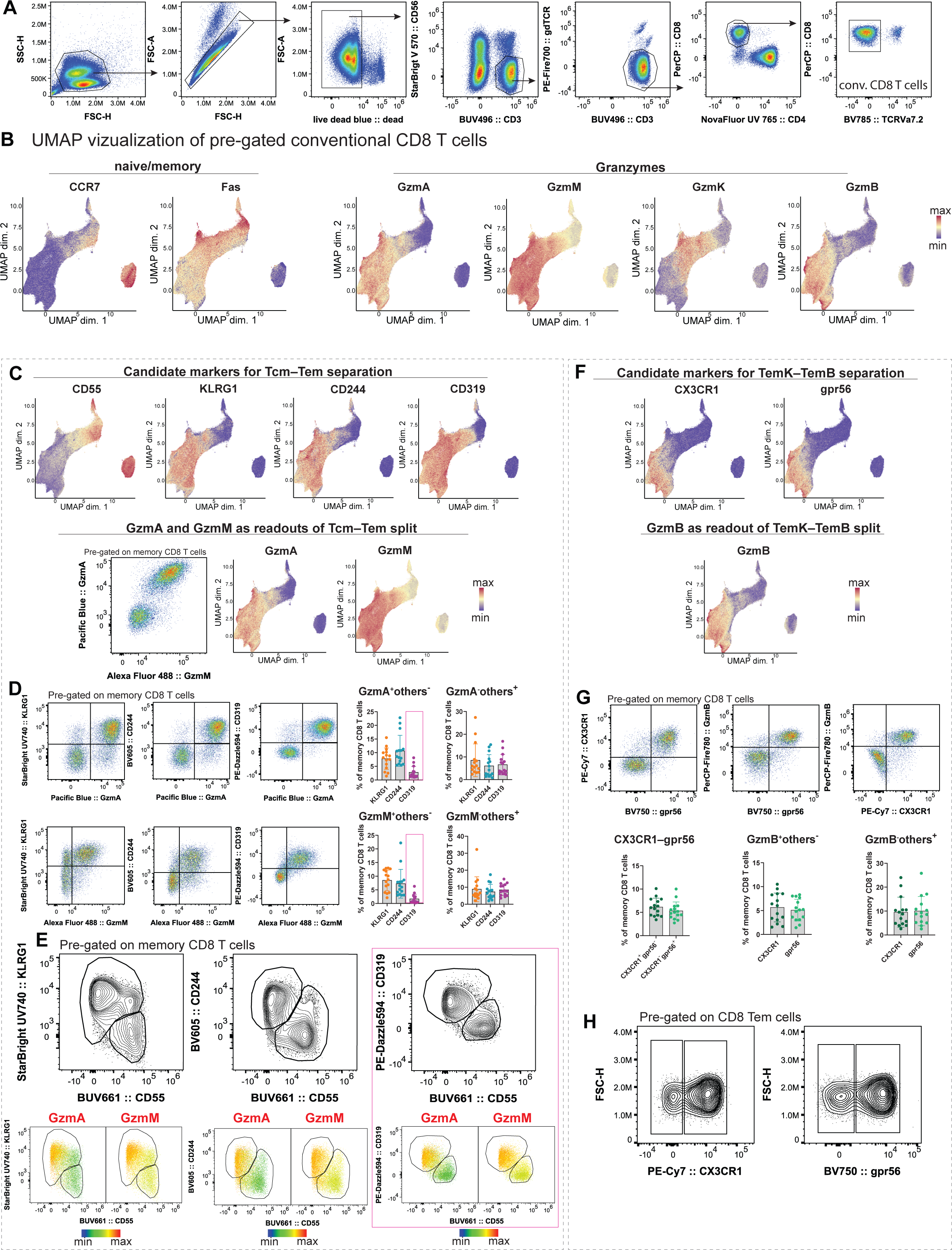
Development of a surface marker–based gating strategy for major CD8 T cell clusters related to Figure 2. (A) Representative gating strategy identifying conventional CD8 T cells. (B) UMAP plots with a protein expression of selected molecules in flow cytometry data. (C) UMAP plots showing expression of candidate surface markers for discrimination of Tcm and Tem CD8 T cell subsets in flow cytometry data (top). Representative dot plot showing co-expression of GzmA and GzmM (bottom left) and UMAP plots with a protein expression of GzmA and GzmM (bottom right). (D) Representative flow cytometry dot plots showing co-expression of KLRG1, CD244, and CD319 with GzmA (top left) or GzmM (bottom left). Bar plots quantifying the extent of co-expression between selected surface markers and GzmA (top right) or GzmM (bottom right). (E) Representative contour plots showing co-expression of KLRG1, CD244, and CD319 with CD55 (top). Representative heat maps highlighting regions enriched for GzmA^+^ or GzmM^+^ cells within KLRG1–CD55, CD244–CD55, and CD319–CD55 co-expression plots (bottom). (F) UMAP plots showing expression of candidate surface markers for discrimination of TemK and TemB cells (top) and GzmB (bottom) in flow cytometry data (G) Representative flow cytometry dot plots showing co-expression of CX3CR1, gpr56, and GzmB (top). Bar plots quantifying the extent of co-expression between selected surface markers and GzmB (bottom). (H) Representative contour plots showing expression of CX3CR1 and gpr56 in effector memory CD8 T cells

**Supplementary Figure 4:**
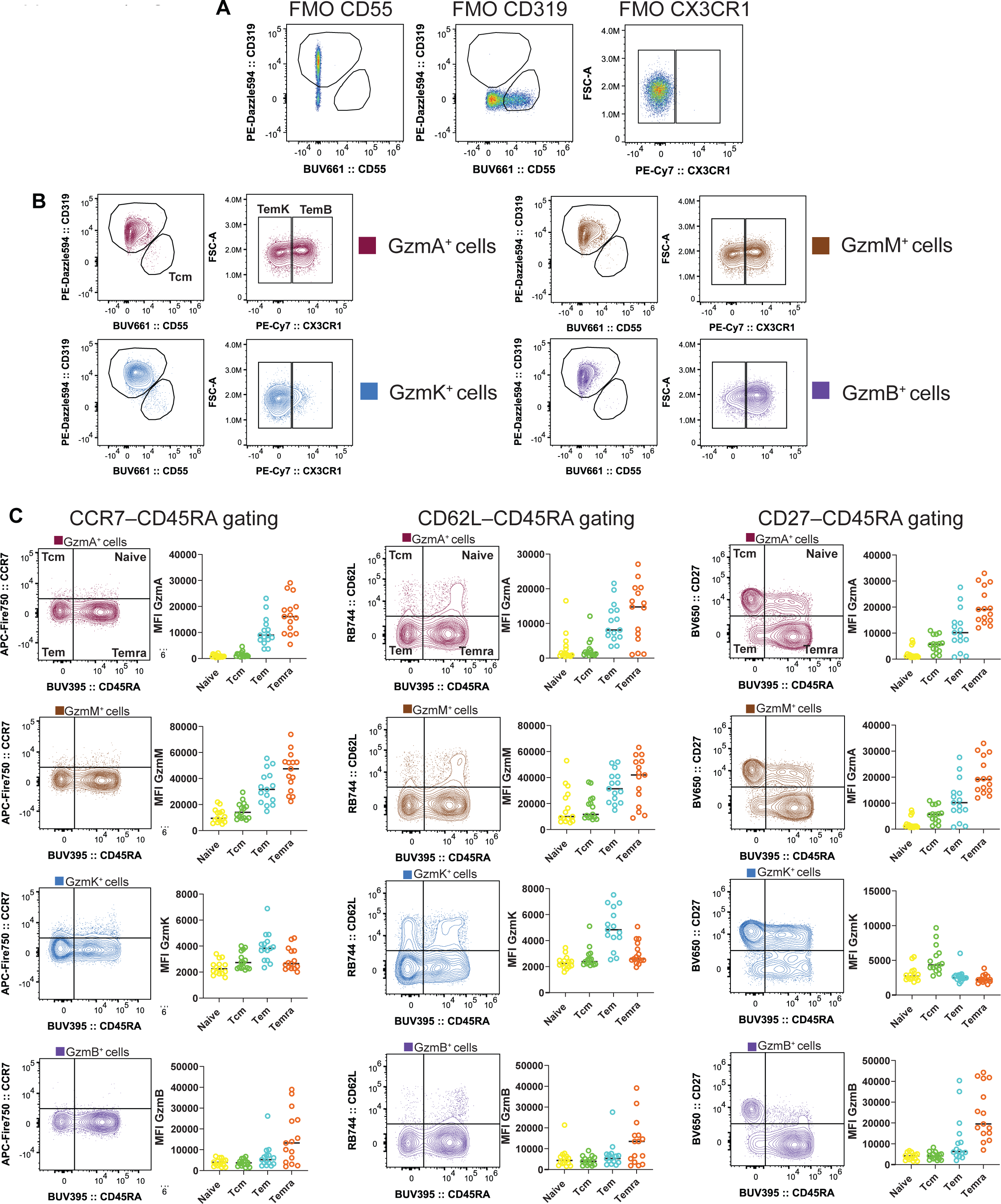
Granzyme expression across CD8 T cell subsets defined by different gating strategies related to Figure 2. (A) Representative flow cytometry dot plots showing FMO controls used for the CD55–CD319–CX3CR1 gating strategy. (B) Representative flow cytometry contour plots showing distribution of GzmA⁺, GzmM⁺, GzmK⁺, and GzmB⁺ CD8 T cells within CD55–CD319–CX3CR1 subsets. (C) Representative flow cytometry contour plots showing the distribution of GzmA⁺, GzmM⁺, GzmK⁺, and GzmB⁺ CD8 T cells within CCR7–CD45RA (left), CD62L–CD45RA (middle), and CD27–CD45RA (right) gating subsets. Adjacent scatter plots show granzyme MFI across the corresponding canonical CD8 T cell subsets.

**Supplementary Figure 5:**
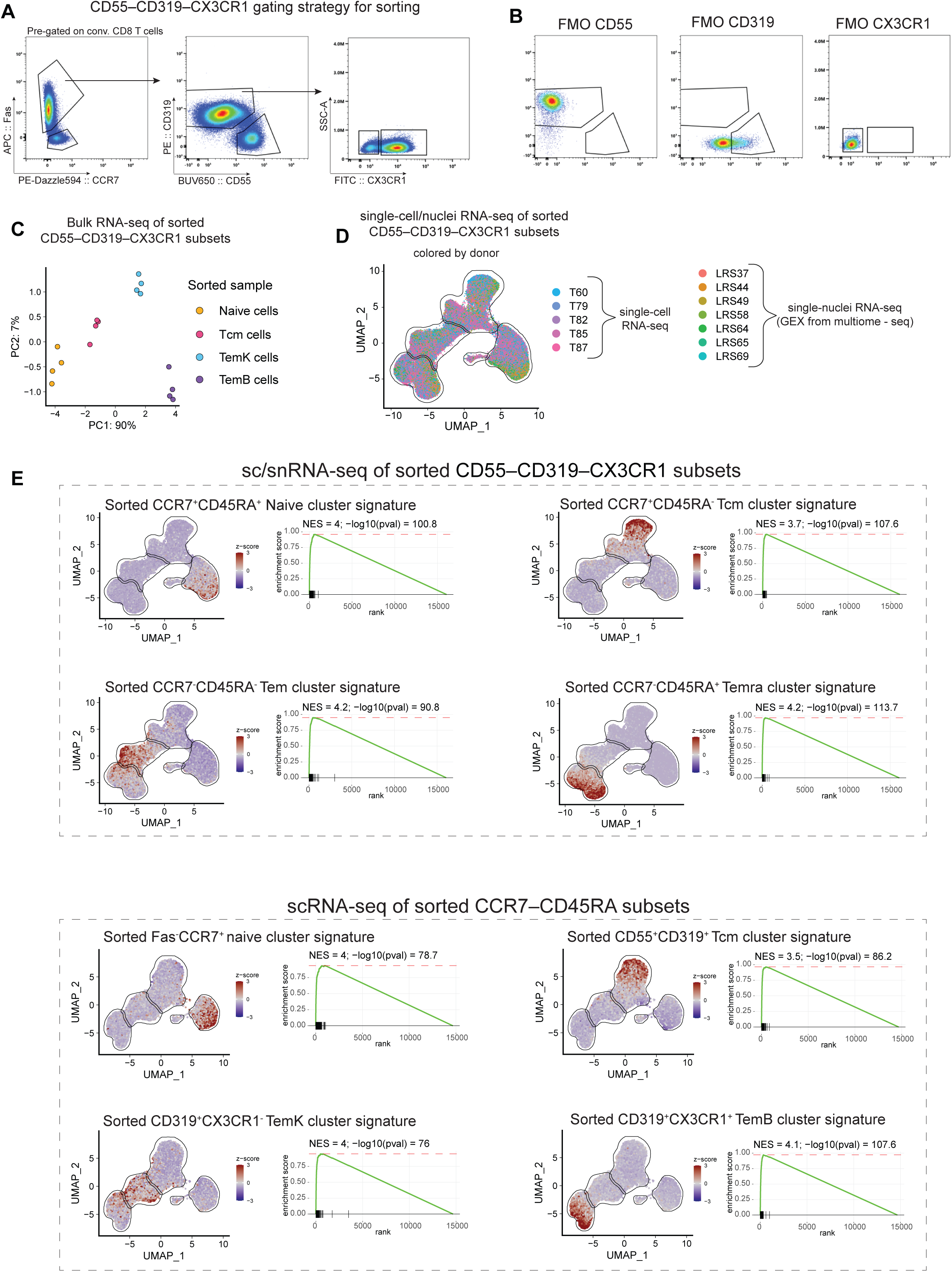
Sorting strategy and transcriptional characterization of CD55–CD319–CX3CR1 subsets with comparison to previously established scRNA-seq of CCR7–CD45RA subsets related to Figure 2. (A) Representative dot plots showing the sorting strategy for CD55–CD319–CX3CR1 subsets. (B) Representative flow cytometry dot plots showing FMO controls used for the CD55–CD319– CX3CR1 sorting strategy. (C) Principal component analysis (PCA) of bulk RNA-seq data from sorted CD55–CD319– CX3CR1 subsets. (D) UMAP plot of CD55–CD319–CX3CR1 subsets, colored by donor. (E) UMAP plots showing gene set coregulation analysis (GESECA) scores of major CD8 T cell cluster signatures and corresponding GSEA enrichment analyses. Signatures from sorted CCR7– CD45RA subsets were projected onto sc/snRNA-seq data of CD55–CD319–CX3CR1-sorted subsets (top), and vice versa (bottom).

**Supplementary Figure 6:**
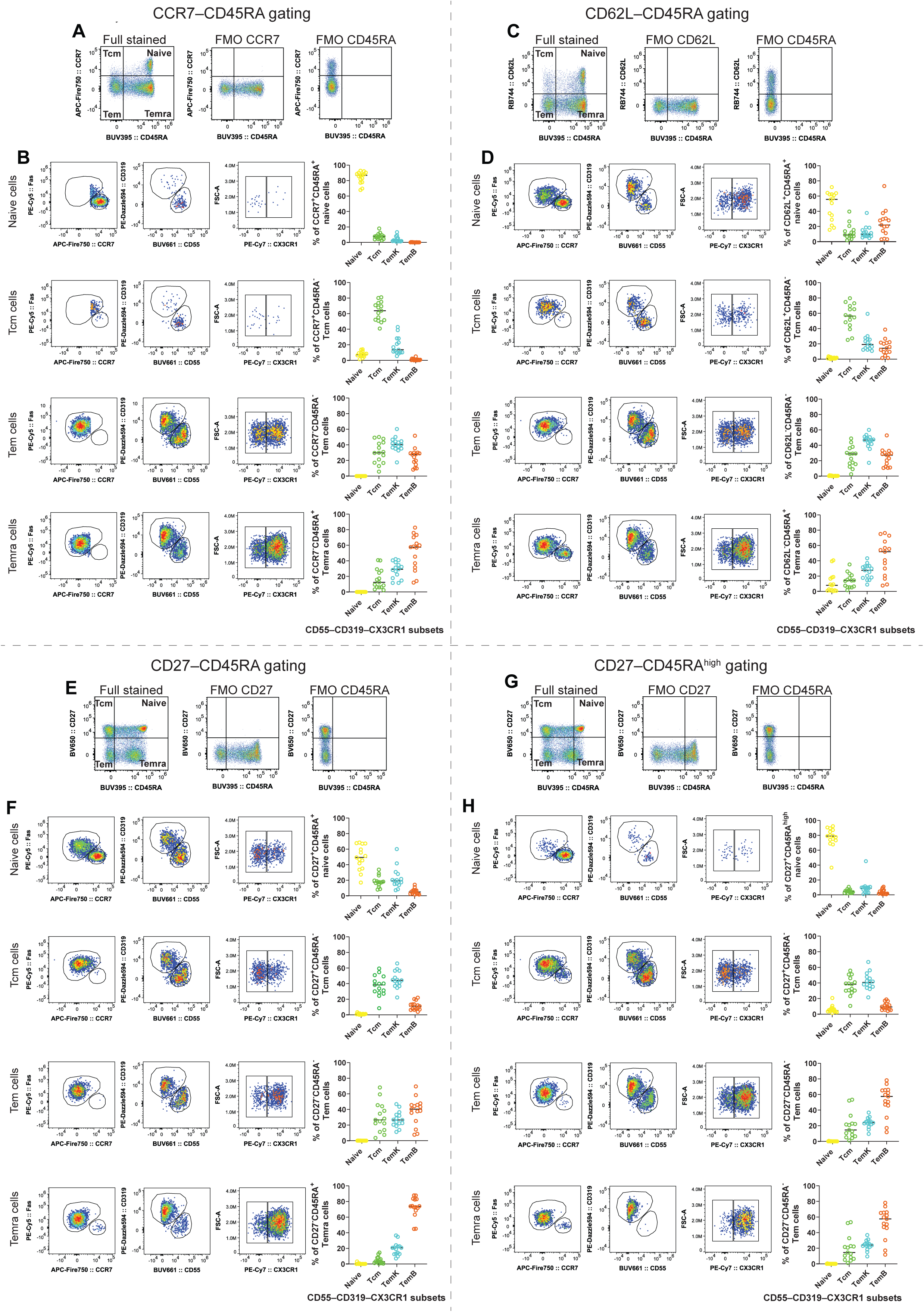
Comparison of canonical CD8 T cell gating strategies and CD55–CD319–CX3CR1-defined subsets related to Figure 2. (A) Representative dot plots showing the gating strategy for CCR7–CD45RA subsets and corresponding FMO controls. (B) Representative flow cytometry dot plots illustrating the distribution of CCR7–CD45RA subsets within the CD55–CD319–CX3CR1 gating framework (left). Scatter plots show the proportion of CCR7–CD45RA subsets assigned to each CD55–CD319–CX3CR1-defined subset across donors (right). (C) Representative dot plots showing the gating strategy for CD62L–CD45RA subsets and corresponding FMO controls. (D) Representative flow cytometry dot plots illustrating the distribution of CD62L–CD45RA subsets within the CD55–CD319–CX3CR1 gating framework (left). Scatter plots show the proportion of CD62L–CD45RA subsets assigned to each CD55–CD319–CX3CR1-defined subset across donors (right). (E) Representative dot plots showing the gating strategy for CD27–CD45RA subsets and corresponding FMO controls. (F) Representative flow cytometry dot plots illustrating the distribution of CD27–CD45RA subsets within the CD55–CD319–CX3CR1 gating framework (left). Scatter plots show the proportion of CD27–CD45RA subsets assigned to each CD55–CD319–CX3CR1-defined subset across donors (right). (G) Representative dot plots showing the gating strategy for CD27–CD45RA^high^ subsets and corresponding FMO controls. (H) Representative flow cytometry dot plots illustrating the distribution of CD27–CD45RA^high^ subsets within the CD55–CD319–CX3CR1 gating framework (left). Scatter plots show the proportion of CD27–CD45RA^high^ subsets assigned to each CD55–CD319–CX3CR1-defined subset across donors (right).

**Supplementary Figure 7:**
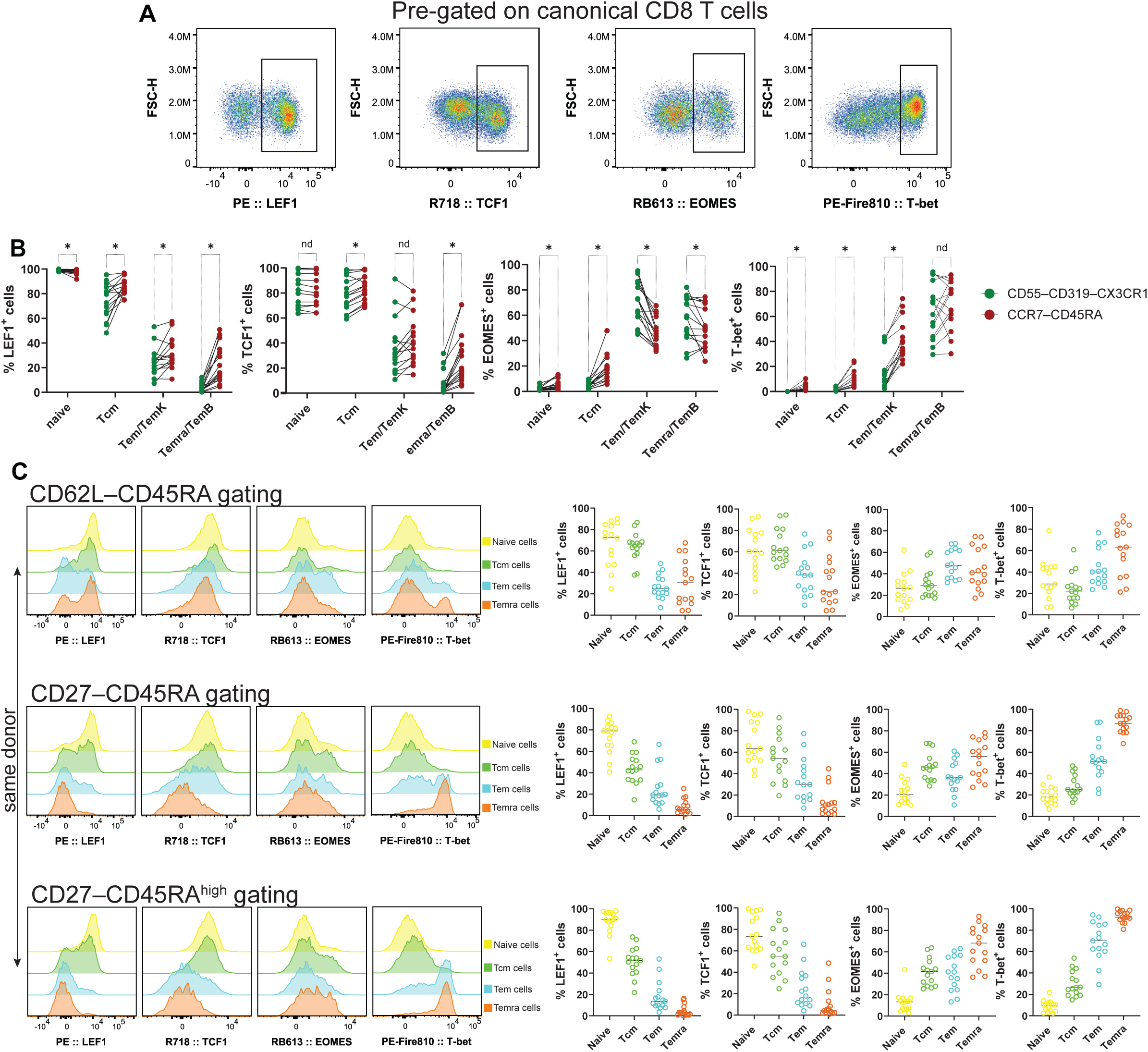
Transcription factor expression across CD8 T cell subsets defined by different gating strategies related to Figure 3. (A) Representative flow cytometry dot plots showing expression of LEF1, TCF1, EOMES, and T-bet within conventional CD8 T cells. (B) Quantification of LEF1⁺, TCF1⁺, EOMES⁺, and T-bet⁺ cells within corresponding subsets defined by CD55–CD319–CX3CR1 (green) or CCR7–CD45RA (red) gating strategies. Statistical significance was assessed using multiple t-tests with FDR correction. Asterisks denote significant discoveries (q < 0.05); nd indicates no significant difference. (C) Representative histograms showing protein transcription factor expression by CD62L– CD45RA, CD27–CD45RA or CD27–CD45RA^high^ subsets (left). Scatter plots showing the percentage of transcription factor positive cells CD62L–CD45RA, CD27–CD45RA or CD27– CD45RA^high^ subsets (right).

**Supplementary Figure 8:**
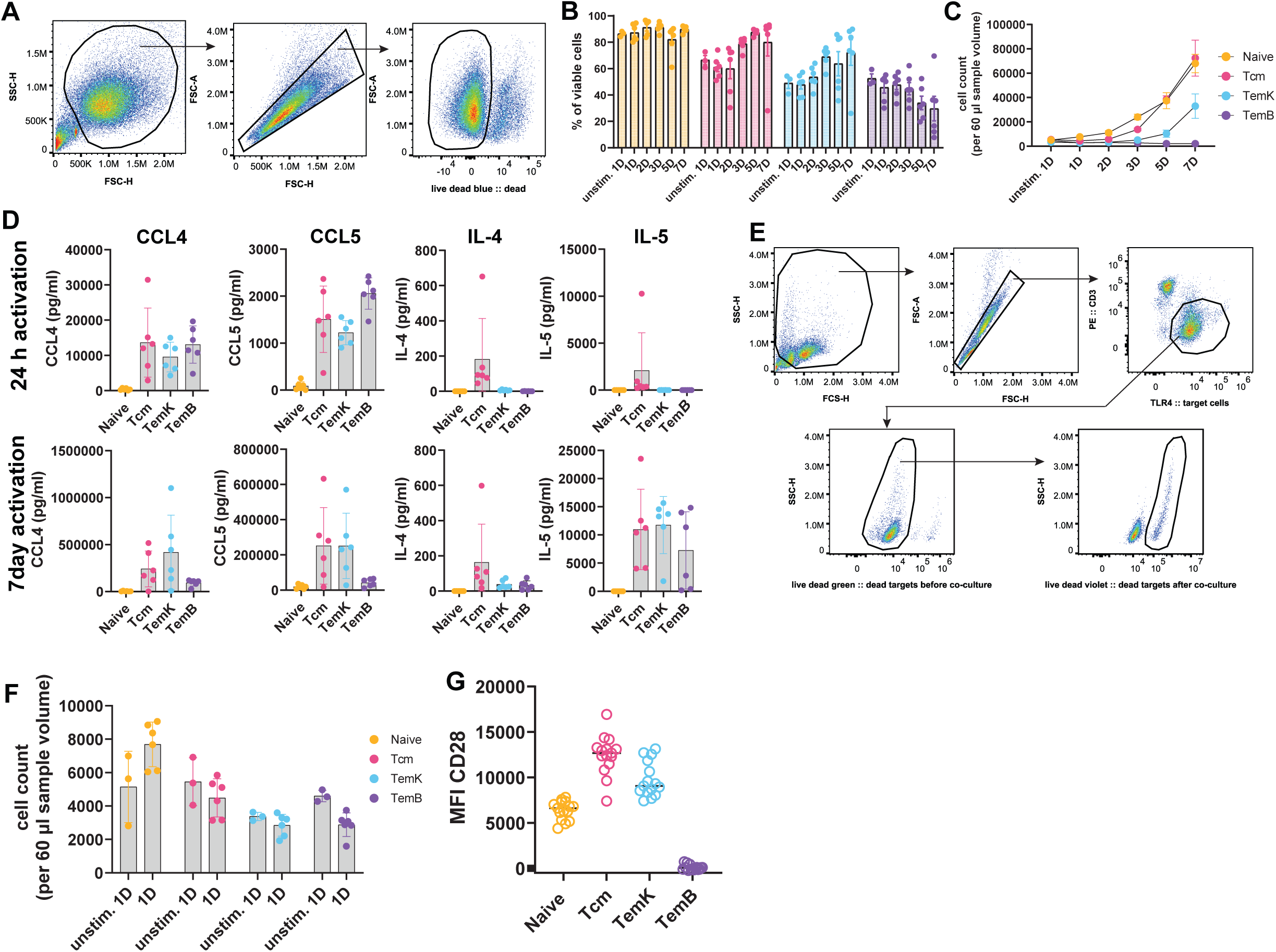
Expanded functional profiling of CD55–CD319–CX3CR1-defined subsets related to Figure 4. (A) Representative gating strategy for αCD3/αCD28 beads activated CD8 T cells. (B) Bar plots showing viability of activated CD8 T cell subsets during time course. (C) Line plots showing cell expansion kinetics of sorted CD55–CD319–CX3CR1 subsets, linear scale. (D) Bar plots showing cytokine production by CD55–CD319–CX3CR1 subsets following 24 hours or 7 days of αCD3/αCD28 bead stimulation. (E) Representative flow cytometry gating strategy used to assess target P815 cell death. (F) Bar plots showing cell counts of unstimulated and stimulated CD55–CD319–CX3CR1 subsets after 24 hours following *in vitro* activation. (G) Scatter plot showing MFI of CD28 in CD55–CD319–CX3CR1 subsets across donors.

## METHODS

### Human samples

The Washington University in St. Louis School of Medicine Institutional Review Board reviewed the collection of blood samples from healthy participants (IRB-201804084). Non-obese (BMI under 30) males and females were enrolled in the study. Participants were given a screening questionnaire to establish their health status. Non-obese (BMI under 30) and non-smoking participants without a history of cancer, chronic inflammatory conditions (arthritis, Crohn’s disease, colitis, dermatitis, fibromyalgia, or lupus), or blood-borne infections (HIV, hepatitis B, and C) were included. Subjects who reported cold or flu symptoms in the prior month were excluded. Peripheral blood (∼ 100 ml) was collected into sodium-heparin BD vacutainer tubes by venous puncture in the morning (7-10 AM) after an overnight fast. Additionally, the leukoreduction filters after platelet apheresis of anonymous healthy blood donors of both sexes were obtained.

### Sample collection

PBMCs were isolated from whole blood by density gradient centrifugation using Histopaque-1077 (Sigma, cat. #10771) according to the manufacturer’s instructions. Briefly, the whole blood was diluted in a 1:1 ratio with DPBS (Gipco, cat. #14190136) with 2mM EDTA (Corning, cat. #46-034-CI). The diluted blood was overlaid on Histopaque-1077 and centrifuged at 500g for 30 min at room temperature without breaks. PBMC were collected from diluted plasma -Histopaque interface and washed twice in DPBS–2mM EDTA. PBMC were cryopreserved in CryoStor CS10 freezing medium (Biolife Solutions, cat. #210502) and stored at –80°C.

### Single-cell RNA-seq of CD8 T cell subsets

Cryopreserved PBMCs were thawed by serial dilution with an equal volume of 2% FBS in PBS. CD8 T cells were enriched from PBMCs by negative selection using magnetic beads (StemCell Technologies, cat. #17953). Cells were stained with Live/Dead Fixable Blue Dead Cell Stain Kit (Invitrogen, cat. #L34962), followed by incubation with Human TruStain FcX blocking solution (BioLegend, cat. #422302) at 4 °C. Cells were then stained with antibody cocktails targeting specific CD8 T cell subsets, along with 0.2 µg of a TotalSeq-C Hashtag antibody per donor (**Table2**). Samples were incubated at 4 °C for 30 minutes. Labeled cells were subsequently sorted into CD8 T cell subpopulations (canonical CCR7–CD45RA subsets **Fig.S1B**; CD55–CD319– CX3CR1-defined subsets **Fig. S5A, B**) using an Aurora Cell Sorter (Cytek Biosciences). The same cell populations from each donor were sorted into the same 1.5 mL tube to generate equal pools. Pooled cell samples were filtered through a 70 µm FloMi filter (SP Bel-Art, cat. #136800070), counted, and assessed for viability using AO/PI staining on a Cellometer™ Ascend (Revvity). The final cell suspension was adjusted to approximately 1,750 cells/µL. scRNA-seq and feature barcode libraries were constructed and sequenced at the McDonnell Genome Institute (Washington University in St. Louis) using Chromium GEM-X Single Cell 5′ Reagent Kits v3 and Illumina NovaSeq X Plus 10B flow cells, targeting 50,000 reads per cell for gene expression and 5,000 reads per cell for feature barcode libraries.

### Bulk RNA seq of CD8 T cell subsets

Cryopreserved PBMCs were thawed and CD8 T cells were enriched by negative selection using magnetic beads (StemCell Technologies, cat. #17953). CD8 T cell subpopulations (CD55– CD319–CX3CR1-defined subsets **Fig. S5A, B**) were sorted using Aurora Cell Sorter (Cytek Biosciences). mRNA was isolated directly from cell pellets with RNeasy Plus Micro Kit (QIAGEN, cat. #74034). RNA-seq libraries were constructed and sequenced at the McDonnell Genome Institute (WASHU, St Louis) using the Takara-Clontech SMARTer system and Illumina NovaSeq6000 S4 XP flow cells with 2x150 paired-end reads.

### Multiome-seq of CD8 T cell subsets

Cryopreserved PBMCs were thawed and CD8 T cells were enriched by negative selection using magnetic beads (StemCell Technologies, cat. #17953). CD55–CD319–CX3CR1-defined subsets of CD8 T cells (**Fig. S5A, B**) were sorted using Aurora Cell Sorter (Cytek Biosciences), and cells were pooled equally into one subtype sample. Permeabilized nuclei for multiome-seq were prepared from the sorted subsets according to the Nuclei Isolation for Single Cell Multiome ATAC + Gene Expression Sequencing protocol for PBMCs (10x Genomics). Nuclei were counted and assessed for integrity using a Cellometer™ Ascend (Revvity) prior to transposition. Single-nuclei multiome libraries were generated using the Chromium Single Cell Multiome ATAC + Gene Expression kit v1.0 (10x Genomics) according to the manufacturer’s instructions. Transposed nuclei were then loaded onto a Chromium X instrument using Chip J (10x Genomics), targeting ∼10,000 nuclei per reaction for GEM generation and barcoding. Gene expression and chromatin accessibility libraries were subsequently prepared from the same nuclei, and library quality and concentration were assessed prior to sequencing. Sequencing was performed on an Illumina NovaSeq X Plus 25B (Illumina) using paired-end reads following 10x Genomics recommendations, targeting ∼80,000 read pairs per nucleus for gene expression and ∼80,000 for ATAC.

### Spectral flow cytometry

Cryopreserved PBMC were thawed and stained with Live/dead fixable blue stain kit (Invitrogen, cat. #L34962), followed by human TruStain FcX blocking solution (Biolegend, cat. #422302) for 15 mins at 4 C. According to previous reports^41,42^, chemokine receptors, γδTCR or other surface markers show decreased resolution between positive and negative signals in multicolor panels potentially due to steric hindrance, so we use the sequential staining protocol to improve the resolution for these markers as we described previously^13,43^. PBMC were pre-stained with a cocktail of resolution-sensitive markers for 5 mins, followed by a direct addition of surface antibody cocktail, and incubated for an additional 30 min at 4 °C. For intracellular staining, Foxp3 / Transcription Factor Staining Buffer Set (eBioscience, cat. #00-5523-00) was used according to the manufacturer’s instruction. The used antibodies are specified in **Table3**. Flow cytometry data were acquired using Cytek Aurora cytometer (Cytek Biosciences) equipped with 5 lasers, followed by spectral unmixing using SpectroFlo software (Cytek Biosciences).

### Stimulation of CD8 T cell subsets

Cryopreserved PBMCs were thawed, and CD8 T cells were enriched by negative selection using magnetic beads (StemCell Technologies, cat. #17953). Enriched cells were sorted into CD8 T cell subpopulations (CD55–CD319–CX3CR1-defined subsets **Fig. S5A, B**) using an Aurora Cell Sorter (Cytek Biosciences). Sorted cells were cultured in RPMI 1640 medium supplemented with 10% FBS, 1% penicillin/streptomycin, and 2 mM L-glutamine at a concentration of 0.3 × 10⁵ cells/mL in a V-bottom 96-well plate. Sorted CD8 T cell subsets were stimulated for 1–7 days with anti-CD3 and anti-CD28 Dynabeads (bead-to-cell ratio 1:2; Gibco, cat. #11161D), supplemented with IL-2 (10 ng/mL, BioLegend, cat. #589102) or IL-7 (10 ng/mL, BioLegend, cat. #581902) and IL-15 (10 ng/mL, BioLegend, cat. #570302). On day 4, 50 µL of fresh medium was added to cultures lasting ≥4 days.

Cell supernatants were collected and analyzed for cytokine levels using multiplex fluorescent bead assays (Eve Technologies; HDF15, HDCD8+ assays, and a custom panel including MIP-1β and RANTES). Cells were subsequently analyzed by flow cytometry for viability and activation markers using a Cytek Aurora cytometer (Cytek Biosciences) equipped with 5 lasers, followed by spectral unmixing using SpectroFlo software (Cytek Biosciences).

### Killing assay

Cryopreserved PBMCs were thawed, and CD8 T cells were enriched by negative selection using magnetic beads (StemCell Technologies, cat. #17953). Enriched cells were sorted into CD8 T cell subpopulations (CD55–CD319–CX3CR1-defined subsets **Fig. S5A, B**) using an Aurora Cell Sorter (Cytek Biosciences).

Sorted cells were cultured in RPMI 1640 medium supplemented with 10% FBS, 1% penicillin/streptomycin, and 2 mM L-glutamine at a density of 10,000 or 40,000 cells per well in a V-bottom 96-well plate. CD8 T cell subsets were stimulated for 24 hours with anti-CD3 and anti-CD28 Dynabeads (bead-to-cell ratio 1:2; Gibco,cat. #11161D), supplemented with IL-2 (10 ng/mL, BioLegend, cat. #589102) or IL-7 (10 ng/mL, BioLegend, cat. #581902) and IL-15 (10 ng/mL, BioLegend, cat. #570302). After stimulation, beads were removed using a magnet, and cytokine-supplemented medium was replaced with fresh complete medium. Cells were rested for at least a 1 hour.

P815 mastocytoma target cells (ATCC, cat. # TIB-64) were labeled with Live/Dead Fixable Green Stain Kit (Invitrogen, #L34969) and TFL4 (OncoImmunin, cat. #PTL802) for 10 minutes in PBS, washed in 20% FBS in PBS, and incubated for 30 minutes at room temperature with α-CD3 (1 µg per million P815 cells; clone OKT3; BD, cat. #571671). A total of 10,000 labeled P815 target cells were added to cultures containing rested CD8 T cell subsets and incubated overnight at 37 °C and 5% CO₂. After incubation, cells were stained with Live/Dead Fixable Violet Stain Kit (Invitrogen, cat. #L34963) and anti-human CD3 antibody conjugated to PE (BioLegend, cat. #344805). Data were acquired using a Cytek Aurora cytometer (Cytek Biosciences) equipped with 5 lasers.

### Computational flow cytometry data processing

PBMC data were manually pre-gated on conventional CD8 T cells (**Fig. S3A**) in FlowJo V 10.10.0 software (Tree Star). Exported CD8 T cells were clustered using Leiden clusterization on 32 markers (CD45RA, HLA DR, CD55, CCR7, CCR9, GzmA, CD45RO, CD244, CD27, integrin B7, CD49d, Cx3CR1, CD49a, CD28, CD29, CD26, GzmK, CD62L, GzmB, CD73, LEF1, CD103, CD319, Fas, HELIOS, KLRG1, CXCR3, CCR10, CD25, TCF1, CCR4, CD38). For

Leiden clusterization, multiple sub-steps were done: (a) k nearest neighbors determining for each event using hnsw_knn function with squared Euclidean parameter from R package RcppHNSW^44^ v0.6.0, (b) calculating nearest-neighbor distances using functions rcpp_parallel_jce and dedup_links from R package FastPG^45^ v0.0.8, (c) Leiden clustering using JAVA library CWTSLeiden/networkanalysis^46^ v1.1.0 and its command-line tool RunNetworkClustering with Modularity and weighted-edges parameters. Parameters k was set to 50 and resolution parameter to 0.185. UMAP construction for the selected markers was created by the UMAP function of R package uwot^47^ v0.2.3 on scaled and centered expression matrix using scale function from R base package^48^ v4.4.0.

### Single-cell data processing

Single-cell data for CCR7-CD45RA and CD55-CD319-CX3CR1 sorting experiments consisted of single-cell gene expression (GEX) and hashtag oligonucleotide (HTO) libraries. For each experiment, each sorted sample was processed independently using the 10x Genomics *Cell Ranger count* pipeline (available at the 10x website). As the data were generated across multiple experiments and processed at different time points, Cell Ranger versions differed between datasets (v10.0.0 and v9.0.1). Following the generation of count matrices, all datasets were subjected to a common downstream analysis pipeline.

Reads were aligned to the human GRCh38 reference transcriptome using refdata-gex-GRCh38-2024-A. For experiments that included HTO libraries, Cell Ranger was run using a library configuration file specifying both GEX and HTO libraries, together with a feature reference file containing the hashtag oligonucleotide barcode sequences and corresponding feature annotations. BAM files were generated during the Cell Ranger run to enable downstream genotype-based demultiplexing.

To resolve pooled donors within each sorted sample, genotype-based demultiplexing was performed using *Souporcell*^49^ v2022.12. For each Cell Ranger output, the aligned BAM file and the corresponding cell barcode file were used as inputs. The number of expected genotype clusters was set according to the number of donors pooled in the corresponding sample. Only cells classified as genotype singlets were retained for downstream analysis.

Final donor identities were assigned by combining genotype-based demultiplexing with HTO signal. Souporcell genotype clusters were first inspected to identify groups of cells corresponding to individual donors. These genotype clusters were then matched to the expected experimental donor identities using the dominant normalized HTO signal detected in each cluster.

For each sorted sample, filtered Cell Ranger matrices were imported into the *R*^48^ environment v4.3.0. Gene expression counts were used to initialize a Seurat object using *Seurat*^50^ package v4.3.0. Each sample was processed independently prior to merging, allowing quality control metrics and donor assignments to be evaluated within each sorted sample. Gene expression counts were normalized using log-normalization with a scale factor of 10,000. Highly variable genes were identified using 1,500 variable features per object. To prevent T-cell receptor genes from driving dimensionality reduction and clustering, genes beginning with TRA or TRB were removed from the variable feature set before scaling and PCA. The data were scaled using the selected variable features. During scaling, total UMI count (nCount_RNA) and mitochondrial percentage (percent.mt) were regressed out using a linear model. This was done to reduce the influence of sequencing depth and cell-quality variation on downstream dimensionality reduction. Principal component analysis was performed on the scaled expression matrix using the selected variable genes. Harmony integration (*harmony*^51^ package v0.1.1) was then applied to the PCA embeddings to correct for donor-associated effects. UMAP dimensionality reduction was performed using the first 15 Harmony dimensions. The same Harmony dimensions were used to construct the nearest-neighbor graph. Graph-based clustering was performed in Seurat using the shared nearest-neighbor graph. Initial sample-level clustering was performed at low resolution to identify major cell populations, technical artifacts, and contaminating clusters.

Quality control was performed at the single-cell level using multiple metrics, including the total number of detected UMIs, the number of detected genes, and the percentage of mitochondrial reads. Cells with low UMI counts or low numbers of detected genes were removed as low-quality cells. Cells with high mitochondrial content were removed to exclude damaged or dying cells. Cells with excessively high numbers of detected genes or UMIs were removed to reduce the contribution of multiplets or other technical artifacts.

In addition to metric-based filtering, a small contaminating γδ T-cell cluster was removed from one of the samples. One donor from the CD55-CD319-CX3CR1 sorting experiment was excluded from downstream analyses because the sample failed quality control and exhibited an unusually high proportion of HLA-DR–expressing cells, indicative of a potentially activated immune state. Following sample-level quality control, demultiplexing, and donor assignment, all filtered single-cell objects belonging to the same sorting experiment were merged into a single Seurat object. This resulted in two independently processed datasets corresponding to the CCR7-CD45RA and CD55-CD319-CX3CR1 sorting experiments. Each merged object was reprocessed using the same general workflow. Gene expression counts were log-normalized with a scale factor of 10,000. Variable features were selected using 1,500 genes, again excluding TCR alpha- and beta-chain genes from the variable feature set. The merged dataset was scaled with a regression of the total UMI count and the mitochondrial percentage.

PCA was performed on the scaled expression matrix, followed by Harmony integration to mitigate donor-associated variation in the merged dataset. Harmony was run using donor identity as the primary integration variable. UMAP visualization was calculated from the Harmony-corrected embeddings using the first 15 dimensions. A nearest-neighbor graph was constructed using the same dimensions, and graph-based clustering was performed using the Louvain algorithm. The final clustering resolution was chosen as the minimum resolution required to resolve the four major CD8 T cell populations (Naive, Tcm, TemK, and TemB). Cluster identities were assigned using established marker genes. Subclusters exhibiting highly similar transcriptional profiles and corresponding to the same biological population were consolidated into a single annotation for downstream analyses.

### Multiome data processing

Single-nuclei multiome files for CD55-CD319-CX3CR1 gating were processed using Cell Ranger ARC (v2.1.0, 10x Genomics) and the GRCh38 reference package (refdata-cellranger-arc-GRCh38-2024-A) to generate paired gene expression and chromatin accessibility count matrices. Genotype-based demultiplexing was performed using *Souporcell*^49^ v2022.12. For three out of four samples, donor assignment was inferred from the gene expression BAM files. One sample (TemK) experienced a wetting error during library preparation, and reliable genotype-based demultiplexing was therefore performed using the chromatin accessibility (ATAC) BAM instead. As HTO labeling was not performed for the multiome experiment, donor correspondence between independently demultiplexed samples was established using genotype concordance. For each Souporcell-assigned donor cluster, donor-specific BAM files were generated and processed through the Souporcell variant-calling workflow to obtain donor-level VCF files containing donor-specific single-nucleotide polymorphism (SNP) genotypes. To identify matching donors across samples, only SNP positions detected in all donor-specific VCF files were retained. Pairwise genotype concordance was then calculated between all donor pairs by determining the fraction of shared SNP positions with identical genotype calls. Donors exhibiting the highest genotype concordance were considered to represent the same individual across samples and were assigned a common donor identifier for downstream analyses. To assign specific donor labels and donor information, the same SNP genotype matching procedure was performed between the multiome dataset and an independent single-cell RNA-sequencing dataset containing the same donors and processed with an HTO library. One donor was removed from the final data because only a small number of cells were recovered after the demultiplexing procedure. Following donor assignment, GEX and ATAC components of the multiome dataset were analyzed independently.

### Gene expression portion of multiome data

For the GEX data, preprocessing was performed using the same workflow described above for the single-cell RNA-sequencing datasets using *Seurat*^50^ package v4.3.0. Cells with high mitochondrial transcript content, low UMI counts, low numbers of detected genes, or unusually high UMI or feature counts were removed. In addition, a small contamination of MAIT cells and proliferating cells was excluded from downstream analyses. After quality control, samples were normalized using log normalization, variable features were identified, and data were scaled while regressing out total UMI counts and mitochondrial transcript percentage. Dimensionality reduction and clustering were performed using PCA, Harmony integration, UMAP visualization, graph-based nearest-neighbor construction, and Louvain clustering, as described for the single-cell RNA-sequencing datasets. Donor identity was used as the integration variable during Harmony correction.

To generate a unified transcriptional dataset for downstream analyses, the final integrated multiome GEX object was combined with the independently generated CD55-CD319-CX3CR1 single-cell RNA-sequencing dataset. Integration was performed using Seurat reciprocal PCA (RPCA)-based anchor integration. Briefly, the combined dataset was first split by donor, and 3,000 highly variable genes were identified within each donor. T-cell receptor genes (TRA, TRB) were excluded from the variable feature set. Integration features were selected across donors, followed by data scaling and PCA within each donor. Integration anchors were identified using the RPCA workflow with the first 30 principal components, and an integrated expression matrix was generated using Seurat’s IntegrateData function. The integrated dataset was subsequently scaled while regressing out total UMI counts and mitochondrial transcript percentage, followed by PCA, UMAP visualization, nearest-neighbor graph construction, and Louvain clustering. The final clustering and annotation were performed as described above for the single-cell RNA-sequencing datasets.

### Chromatin accessibility portion of multiome data

Individually generated BAM files of snATAC-seq bulk post-sorted reads were additionally processed with *Omnipeak*^52^ v1.4.6808 in a dedicated snATAC-seq mode with the following parameters:

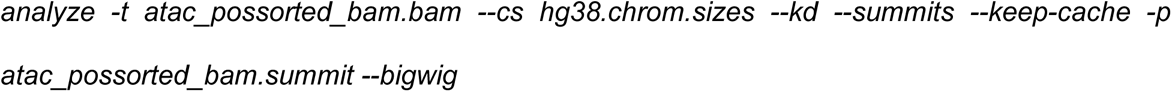

Produced summits were analyzed and compared with Cell Ranger ARC peak-calling results, yielding ∼180k summits per dataset versus ∼100k peaks by Cell Ranger, while maintaining high dataset agreement, yet improving the sensitivity of the downstream analysis. Union peak set was combined using *bedtools*^53^ v2.31.0. Chromatin accessibility data were analyzed using *Signac*^54^ v1.8.0 package. Fragment counts overlapping the union peak set were quantified for each cell using Signac’s *FeatureMatrix* function. A chromatin accessibility assay was constructed using the resulting peak-by-cell count matrix together with the fragment files and gene annotations derived from Ensembl release 86. The chromatin accessibility matrix was normalized using term frequency-inverse document frequency (TF-IDF) transformation. Highly informative peaks were identified using Signac’s *FindTopFeatures* function with a minimum cutoff of the fifth quantile. Dimensionality reduction was performed using latent semantic indexing (LSI) through singular value decomposition. To mitigate donor-specific effects, Harmony integration was applied to LSI dimensions using donor identity as the integration variable. Correlations between sequencing depth and latent dimensions were evaluated using depth-correlation analysis, and dimensions strongly associated with sequencing depth were excluded from downstream analyses. UMAP visualization was generated using Harmony-corrected LSI dimensions 2 and 4-21.

Differential chromatin accessibility analysis was performed between sorted populations using the Wilcoxon rank-sum test as implemented in the Presto package with *min.pct = 0* and *logfc.threshold = 0* parameters. Differentially accessible peaks were annotated to nearby genes using the *ClosestFeature* function implemented in Signac package.

To identify transcription factor binding motifs associated with population-specific chromatin accessibility programs, motif enrichment analysis was performed using *HOMER*^55^ tool v5.1. For each population, the top 1,000 differentially accessible peaks ranked by differential accessibility statistics (average log2 fold change) were selected and exported as BED files. Motif enrichment was then performed using findMotifsGenome.pl against the hg38 reference genome with region-size normalization (-size given).

### Bulk RNA-seq data processing

Bulk RNA-seq samples were processed from paired-end FASTQ files. Raw sequencing read quality was assessed using *FastQC*^56^ v0.11.9, and aggregate quality-control reports were generated using *MultiQC*^57^ v1.34. Paired-end reads were aligned to the human reference genome GRCh38 using *STAR*^58^ v2.7.0f. Alignment quality was assessed using STAR output summaries and visualized across samples using MultiQC. Gene-level read counts were generated using *featureCounts*^59^ v2.0.0. Counts were assigned using the GENCODE v49 primary assembly gene annotation file. Library strandedness parameter (-s 0) for featureCounts was evaluated using *infer_experiment.py* from *RSeQC*^60^ v2.6.4. The resulting gene-by-sample count matrix was used for downstream bulk RNA-seq analysis.

Downstream bulk RNA-seq analyses were performed in the *R*^48^ environment v4.3.0. For visualization and principal component analysis, count matrices were imported and normalized using the *edgeR*^61^ package v3.42.4. Briefly, counts were converted into a DGEList object, library-size normalization factors were calculated, and normalized log-counts per million (logCPM) values were generated using a prior count of 1. Batch correction was performed on logCPM values using *ComBat* function from *sva*^62^ package v3.48.0. Replicate identity was modeled as the batch variable, while the biological sorting population was included in the model matrix to preserve population-specific transcriptional differences during batch correction. PCA was calculated using *prcomp* function from *stats*^48^ package v4.3.0.

Differential expression analysis was performed using *DESeq2*^63^ v1.40.1 on raw count matrices. Prior to model fitting, lowly expressed genes were removed by requiring a minimum of 10 total counts across all samples. Size-factor normalization and dispersion estimation were performed using the DESeq2 workflow, followed by negative binomial generalized linear model fitting and Wald test-based significance testing. Samples were grouped into four sorted populations: Naive, Tcm, TemK, and TemB. For each population, differential expression was tested using a one-versus-rest design. The DESeq2 model included replicate identity as a covariate in the design formula. Differential expression results were generated separately for each population compared with all remaining populations. Log2 fold changes were shrunken using the normal shrinkage estimator. Ranked gene lists (based on Wald test statistic) were used for downstream analyses.

### Publicly available single-cell data

Terekhova et al.^13^ ABF300 CD8 T-cell data were obtained from Synapse: syn49637038. Stuart et al.^64^ Azimuth reference dataset was downloaded from https://atlas.fredhutch.org/nygc/multimodal-pbmc/, and the provided H5Seurat object was loaded using the *SeuratDisk*^65^ v0.0.0.9020 *LoadH5Seurat* function. Data from Gong et al.^66^ were obtained from https://apps.allenimmunology.org/aifi/resources/imm-health-atlas/, and loaded into Seurat-object via *open_matrix_anndata_hdf5* function from *BPCells*^67^ package v0.3.1. Data from Li et al.^24^ were downloaded from GEO accession GSE193442. MAIT and proliferating cells were removed where applicable. For datasets with available embeddings, the original UMAP coordinates were used. The Li et al. dataset did not include cell annotations or UMAP coordinates. These were reproduced by running the analysis pipeline provided by the authors (https://github.com/davis-lab-stanford/kir-cd8) without modification of any parameters. Author-provided (and reproduced) cell annotations were visualized and consolidated into four major CD8 T-cell populations (Naive, Tcm, TemK, and TemB) based on the expression of canonical marker genes.

### Pseudobulk differential expression and gene set enrichment analysis

To identify genes associated with individual single-cell CD8 T cell populations, a pseudobulk differential expression analysis was performed. Raw UMI counts were aggregated across cells from the same donor and cluster using Seurat’s *AggregateExpression* function, generating one pseudobulk profile per donor-cluster combination. Genes with low expression were filtered using the *filterByExpr* function from the *edgeR*^61^ package v3.42.4. Library sizes were normalized using the trimmed mean of M-values (TMM) method. Differential expression analysis was then performed using the *limma-voom*^68^ workflow v3.56.1. Briefly, normalized count data were transformed using the voom method to estimate the mean-variance relationship and generate precision weights. Linear models were fitted with donor identity included as a covariate. For each cluster, differential expression was assessed by comparing pseudobulk samples from that cluster against all remaining clusters combined. Empirical Bayes moderation of standard errors was performed using the eBayes function. Genes were ranked based on t-statistics.

Gene set enrichment analysis was performed to compare cluster-specific transcriptional signatures between datasets. Based on the ranking, the top 100 genes were selected as the cluster-specific gene set. These gene sets were then tested for enrichment against ranked differential expression signatures using *fgseaMultilevel* function from the *fgsea*^69^ package v1.26.0. GESECA scores and corresponding UMAP visualizations were performed using fgsea69 package v1.26.0.

### Estimation of sc/snRNA-seq cluster capture by flow cytometry

To quantify the capture percentage of sc/snRNA-seq clusters by a specific flow cytometry gating strategy, we first calculated cluster frequencies within each sorted sample for every donor. The proportion of each cluster (*Xij*) within a sorted population was determined by dividing the number of cells assigned to that cluster by the total number of cells in the corresponding sorted sample. These cluster proportions were then weighted by the experimentally measured percentage of each sorted population obtained from flow cytometry. Specifically, for each donor, the contribution of a given sorted population to a cluster (*Yij*) was calculated as the product of the cluster proportion (*Xij*) within the sorted sample and the percentage of the corresponding sorted population measured by flow cytometry (*Sj*). Contributions were summed across all sorted populations for each cluster (*∑j Yij*). Finally, the specific capture of a cluster within the sample *(Cij)* was calculated by dividing the weighted contribution of a sorted population to a cluster (*Yij*) by the total contribution of all sorted populations to that cluster (*∑j Yij*) and expressing the result as a percentage.

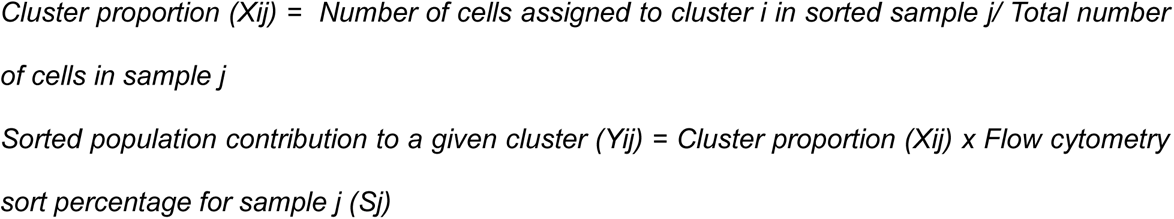

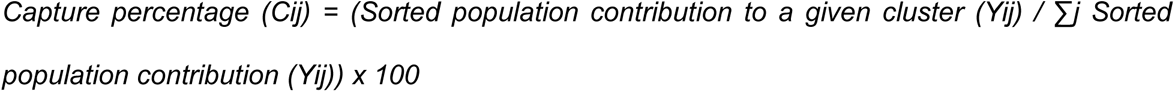

### Visualization

Bar plots, pie charts, box plots, scatter plots, density plots, and enrichment plots were generated using *ggplot2*^70^ v3.4.2. UMAP projections and violin plots were generated using *Seurat*^50^ v4.3.0. Rasterization of high-density single-cell plots was performed using *ggrastr*^71^ v1.0.1. Heatmaps were generated using *pheatmap*^72^ v1.0.12. Genome browser-style coverage tracks were generated using *Signac*^54^ v1.8.0. UMAP density plots were visualized using *ggpointdensity*^73^ v0.2.0. Cell population borders on UMAP embeddings were visualized using *mascarade*^74^ v0.3.1. Peak overlaps between CD8 T cell sorted samples were visualized using a custom UpSet-style plot. Significant (p adjusted value < 1 x 10^-5^ & log fold change > 0) peaks were assigned to population-specific and pairwise intersections and were visualized using *ggplot2*^70^ v3.4.2 and *patchwork*^75^ v1.1.2.

## Data availability

Raw and processed scRNA-seq, multiome-seq data, bulk RNA-seq data generated in this study have been deposited Synapse repository (syn75396198) and are publicly available as of the date of publication. Links for the single-cell online browser have also been deposited at the Synapse repository (syn75396198). Any additional information required to reanalyze the data reported in this paper is available from the lead contact upon request.

